# Adaptive evolution targets a piRNA precursor transcription network

**DOI:** 10.1101/678227

**Authors:** Swapnil S. Parhad, Tianxiong Yu, Gen Zhang, Nicholas P. Rice, Zhiping Weng, William E. Theurkauf

## Abstract

In *Drosophila*, transposon-silencing piRNAs are derived from heterochromatic clusters and a subset of euchromatic transposon insertions, which are transcribed from internal non-canonical initiation sites and flanking canonical promoters. Rhino binds to Deadlock, which recruits TRF2 to promote non-canonical transcription of these loci. Cuff co-localizes with Rhino and Del. The role of Cuff is less well understood, but the *cuff* gene shows hallmarks of adaptive evolution, which frequently targets functional interactions within host defense systems. We show that *Drosophila simulans cuff* is a dominant negative allele when expressed in *Drosophila melanogaster*, where it traps Deadlock, TRF2 and the transcriptional co-repressor CtBP in stable nuclear complexes. Cuff promotes Rhino and Deadlock localization, driving non-canonical transcription. CtBP, by contrast, suppresses canonical cluster and transposon transcription, which interferes with downstream non-canonical transcription and piRNA production. Cuff, TRF2 and CtBP thus form a network that balances canonical and non-canonical piRNA precursor transcription.

## INTRODUCTION

Transposable elements (TEs) are major genome components that can induce mutations and facilitate ectopic recombination, but transposons have also been co-opted for normal cellular functions, and transposon mobilization has rewired transcriptional networks to drive evolution (Ayarpadikannan and Kim, 2014; Hedges and Deininger, 2007; Horvath et al., 2017; Piacentini et al., 2014); (Chuong et al., 2017; Jangam et al., 2017). Species survival may therefore depend on a balance of transposon silencing and activation. The Piwi interacting RNA (piRNA) pathway transcriptionally and post-transcriptionally silences transposons in the germline (Biemont and Vieira, 2006; Canapa et al., 2015; Ghildiyal and Zamore, 2009; Parhad and Theurkauf, 2019; Weick and Miska, 2014). However, how this pathway is regulated is not completely understood.

In *Drosophila*, piRNAs are produced from heterochromatic clusters composed of complex arrays of nested transposon fragments, which appear to provide genetic memory of past genome invaders (Bergman et al., 2006; Brennecke et al., 2007). Adaptation is proposed to require transposition of the new invaders into a cluster, which leads to sequence incorporation into precursors that are processed into trans-silencing anti-sense piRNAs (Khurana et al., 2011). However, a subset of isolated transposon insertions also produce sense and anti-sense piRNAs (Mohn et al., 2014). Formation of these “mini-clusters”, through an uncharacterized epigenetic mechanism, provides an independent adaptation mechanism. In *Drosophila*, germline piRNA clusters and transposon mini-clusters are bound by the RDC complex, which is composed of the HP1 homolog <u>R</u>hino (Rhi), which co-localizes with the linker protein <u>D</u>eadlock (Del) and the Rai1 homolog <u>C</u>utoff (Cuff) (Chang et al., 2019; Chen et al., 2016; Le Thomas et al., 2014; Mohn et al., 2014; Pane et al., 2011; Parhad et al., 2017; Yu et al., 2015; Zhang et al., 2018; Zhang et al., 2014). The three components of the RDC are co-dependent for localization to clusters, and essential to germline piRNA production. Rhino is composed of chromo, hinge and shadow domains (Vermaak et al., 2005). The chromo domain binds to nucleosomes with the H3K9me3 histone modification, and the shadow domain binds Deadlock, which recruits Moonshiner and TATA box related protein 2 (TRF2), promoting “non-canonical” transcription from both genomic strands (Andersen et al., 2017; Le Thomas et al., 2014; Mohn et al., 2014). Mutations in *cuff* trigger a collapse in germline piRNA production, loss of Rhino and Deadlock localization, and a significant increase in cluster transcript splicing (Pane et al., 2011; Zhang et al., 2014), while tethering Cuff to a reporter transgene transcript increases read-through transcription (Chen et al., 2016). The molecular basis for these phenotypes has not been established.

All three RDC genes are rapidly evolving under positive selection, suggesting that the complex is engaged in a genetic conflict (Blumenstiel et al., 2016; Lee and Langley, 2012; Parhad and Theurkauf, 2019; Simkin et al., 2013). We previously found that this process has modified the Rhino-Deadlock interface, producing orthologs that function as mutant alleles when moved across species (Parhad et al., 2017; Yu et al., 2018). Analysis of these “alleles” defined an interaction between the Rhi shadow domain and Del that prevents ectopic assembly of piRNA cluster chromatin. Cross-species analysis of rapidly evolving genes thus offers a potentially powerful genetic approach to structure-function analysis. Here we apply this approach to *cuff*. These studies indicate that adaptive evolution has targeted interactions between Cuff and the Del-TRF2 non-canonical transcriptional complex, and the transcriptional co-repressor <u>C</u>-terminal <u>B</u>inding <u>P</u>rotein (CtBP). CtBP was first identified as a host binding partner of Adenovirus E1A, and subsequently implicated in diverse developmental pathways and cancer (Boyd et al., 1993; Chinnadurai, 2002; Dcona et al., 2017; Mani-Telang et al., 2007; Schaeper et al., 1995; Stankiewicz et al., 2014). We show that *Drosophila* CtBP suppresses canonical transcription from transposon promoters, and promoters flanking two major germline piRNA clusters. Significantly, activation of canonical transcription interferes with downstream non-canonical transcription and piRNA production. Adaptive evolution has therefore targeted interactions between Cuff and two highly conserved transcription regulators, which coordinately regulate germline piRNA expression.

## RESULTS

### *D. simulans cuff* is a dominant separation of function allele in *D. melanogaster*

The *cuff* gene shows hallmarks of adaptive evolution, which can target critical interactions within host defense systems. To determine the functional consequences of *cuff* evolution, we expressed GFP tagged *D. simulans* Cuff (*sim-*Cuff) and control GFP tagged *D. melanogaster* Cuff *(mel-*Cuff) in *D. melanogaster cuff* mutants, and assayed for phenotypic rescue. Both Cuff variants were expressed using the germline-specific *rhi* promoter and integrated into the same chromosomal location, using PhiC31 mediated transformation (Figure 1A). Direct visualization of GFP signal and proteomic studies, described below, indicate that *sim-*Cuff and *mel-*Cuff are nuclear and expressed at comparable levels. Mutations in *cuff* lead to female sterility and production of eggs with dorsal appendage defects, which reflect disruption of D-V patterning in response to genome instability (Klattenhoff et al., 2007). The *mel-cuff* transgene restored D-V patterning and hatching, but the *sim-cuff* transgene failed to rescue either, and was comparable to the null allelic combination in these biological assays. (Figure 1B).

**Figure 1:**
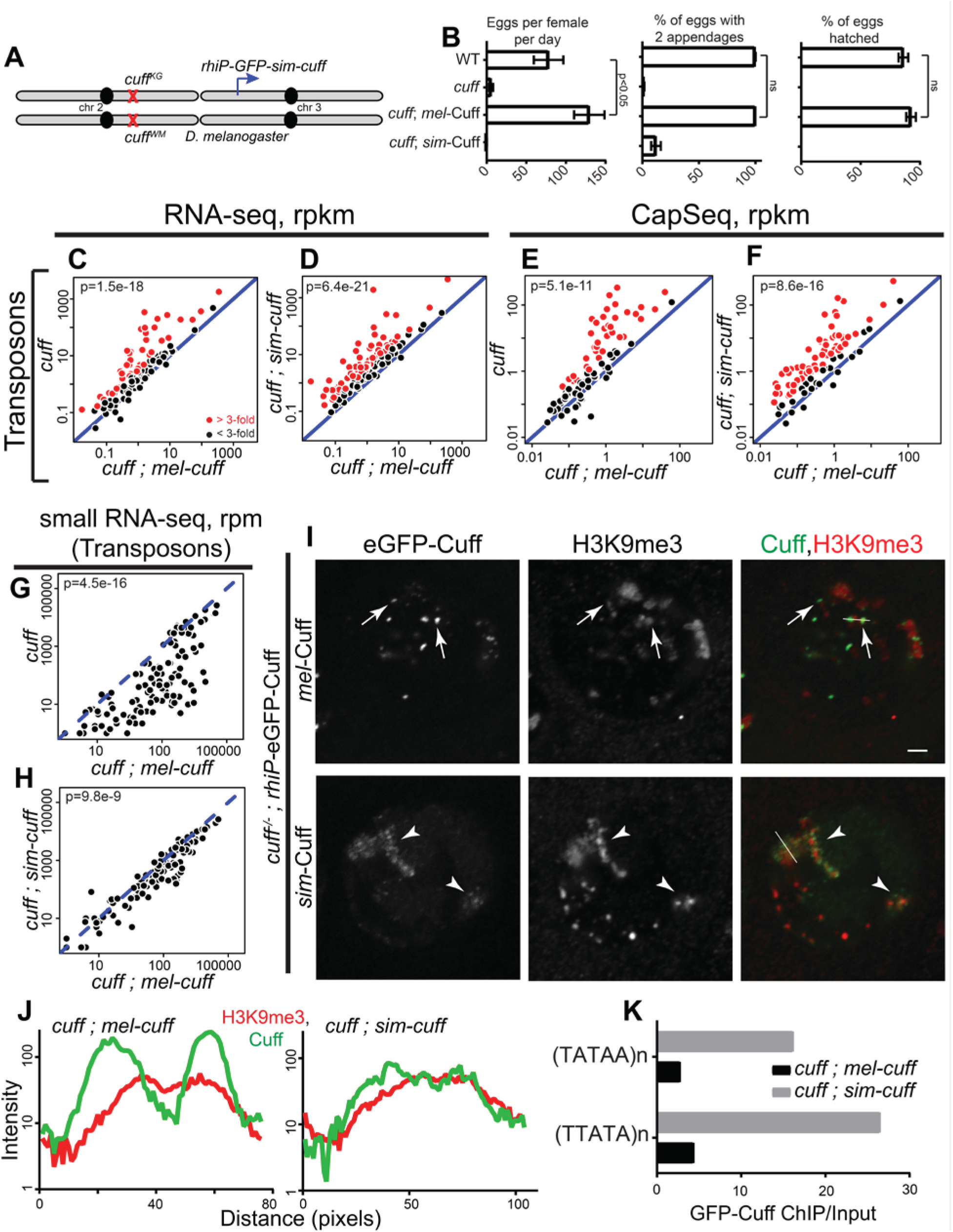
*sim-*cuff does not complement *D. melanogaster cuff* mutations. (A) Genetic complementation strategy. The *sim-cuff* gene was expressed in *D. melanogaster cuff* mutants using the germline specific *rhi* promoter. Similar expression of *mel-cuff* was used as a control. (B) Bar graphs showing number of eggs laid per female per day, % of eggs with 2 appendages and % of hatched eggs produced by OrR (wild type (WT) control), *cuff* mutants, and *cuff* mutants expressing either *mel*-*cuff* or *sim*-*cuff*. Error bars show standard deviation of three biological replicates, with a minimum of 500 embryos scored per replicate, except for *cuff* mutants and *cuff* mutants rescued by *sim-cuff*, where average of 230 and 23 eggs were scored respectively. (C-H) Scatterplots showing comparisons of RNA-seq signal (C, D), CapSeq signal (E, F) and small RNA-seq signal (G, H) at transposons in *cuff* mutant or *cuff* mutant expressing *sim-cuff* vs. *cuff* mutant expressing *mel-cuff*. Each point on the scatterplots shows rpkm (long RNAs) or rpm (small RNAs) for a transposons family in ovaries of the indicated genotype. Diagonal represents x=y. Points in red show y/x>3. p value for differences obtained by Wilcoxon test. (I, J) Localization of GFP tagged Cuff with respect to H3K9me3 marked chromatin in germline nuclei of *cuff* mutants expressing *rhi* promoter driven *mel*-Cuff or *sim*-Cuff. Color assignments for merged images shown on top. Arrowheads and arrows denote locations of *mel*-Cuff or *sim*-Cuff foci respectively. Scale bar, 2 µm. Fluorescence intensities are calculated across the white lines and shown in (J). (K) Bar graphs showing ChIP vs. Input enrichment for GFP-Cuff at the indicated heterochromatic repeats in *cuff* mutant ovaries expressing *mel-cuff* (black) or *sim-cuff* (grey).

To determine if *sim-*Cuff supports transposon silencing (Chen et al., 2007; Pane et al., 2011), we used CapSeq (Gu et al., 2012) and strand-specific RNA-sequencing (Zhang et al., 2012b) to assay steady state expression of transposons and genes. Consistent with the biological assays, the *mel-cuff* transgene restored transposon silencing, and the *sim-cuff* transgene failed to restore silencing (Figures 1C to 1F). Surprisingly, a number of transposon families were more highly expressed in *cuff* mutant expressing *sim-cuff* than in the parental null *cuff* mutant combination (Figures S1A to S1D). Cuff is required for piRNA biogenesis, and small RNA sequencing showed that the *mel-cuff* transgene restored transposon and cluster mapping piRNA expression (Figures 1G and 2D). Based on the transposon silencing defects noted above, we anticipated that *sim-cuff* would also fail to support piRNA expression. However, median transposon and cluster mapping piRNA levels were restored to 45% and 70% respectively, of control levels by the *sim-cuff* transgene, and many clusters and transposons showed essentially wild type piRNA profiles (Figures 1H, S1E, 2E and S2C). Adaptive evolution of *cuff* has therefore generated a *D. simulans* ortholog that functions as a partial separation-of-function allele in *D. melanogaster*, which largely supports piRNA expression, but not transposon silencing.

**Figure 2:**
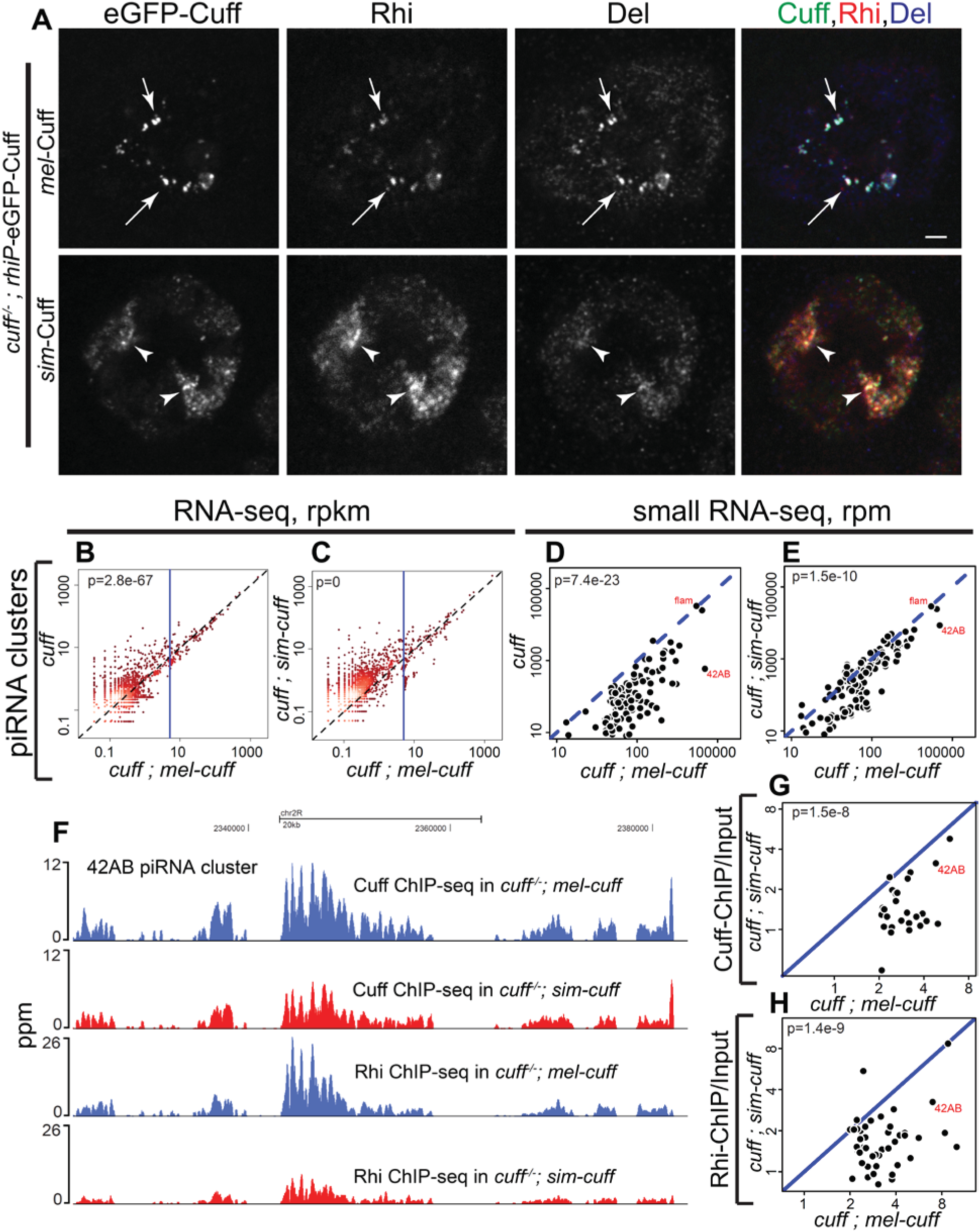
*sim-*Cuff disrupts RDC localization. (A) Localization of GFP tagged Cuff with respect to Rhi and Del in the germline nuclei of *cuff* mutants expressing *rhi* promoter driven *mel*-Cuff or *sim*-Cuff. Color assignments for merged images shown on top. Arrows and arrowheads denote locations of *mel*-Cuff or *sim*-Cuff foci respectively. Scale bar, 2 µm. (B-E) Scatterplots showing comparisons of RNA-seq signal (B, C) and small RNA-seq signal (D, E) at piRNA clusters in ovaries with genotypes *cuff* mutant or *cuff* mutant expressing *sim-cuff* vs. *cuff* mutant expressing *mel-cuff*. In (B, C), each point on the scatterplots shows rpkm value for a 1kb piRNA clusters bin. In (D, E), each point shows rpm value for an entire cluster. Diagonal represents x=y. p value for differences obtained by Wilcoxon test. (F) Genome browser view of GFP-Cuff (top) and Rhi (bottom) ChIP-seq profiles at 42AB piRNA cluster in the ovaries of *cuff* mutants expressing either *mel-cuff* (blue) or *sim-cuff* (red). (G, H) Scatterplots showing comparisons of ChIP/Input values for GFP-Cuff (G) and Rhi (H) at piRNA clusters in ovaries with genotypes *cuff* mutant expressing *sim-cuff* vs. *mel-cuff*. The clusters with prominent Cuff or Rhi binding (rpkm>2) in *cuff* mutant with *mel-cuff* control were used for analysis. Diagonal represents x=y. p value for differences obtained by Wilcoxon test.

Cuff, Rhi and Del associate with peri-centromeric piRNA clusters and localize to cytologically distinct nuclear foci that are frequently adjacent to large domains of constitutive heterochromatin, marked by H3K9me3 (Mohn et al., 2014; Parhad et al., 2017). Consistent with the data presented above, the control *mel-*Cuff:GFP fusion localizes to foci adjacent to these H3K9me3 domains. By contrast, *sim*-Cuff:GFP broadly co-localizes with H3K9me3, and to distinct foci embedded within these domains (Figures 1I and 1J). To determine if *sim-*Cuff disrupts localization of other RDC components, we labeled ovaries expressing the Cuff:GFP fusions for Rhi and Del (Figure 2A). Both proteins colocalized with *mel-*Cuff and *sim-*Cuff, indicating that *sim-* Cuff recruits the RDC to bulk heterochromatin.

To assay RDC localization at the genome level, we performed ChIP-seq for Cuff and Rhino in *cuff* mutants expressing *mel-*Cuff or *sim-*Cuff. As shown in the genome browser view in Figure 2F, *sim-*Cuff fusion shows reduced binding to the 42AB cluster relative to the *mel-*Cuff control, and this is accompanied by reduced Rhi binding. The scatter plots in Figures 2G and 2H compare Cuff and Rhi ChIP-seq enrichment at all clusters, on rescue with *sim-cuff* (y-axis) relative to the *mel-cuff* control (x-axis). Rescue with *sim-cuff* leads to reduced cluster binding by Cuff and Rhi across the genome. Our cytological studies show that *sim-*Cuff leads to RDC co-localization with constitutive heterochromatin (Figures 1I, 1J and 2A). Consistent with these findings, *sim-*Cuff shows enhanced binding to two A/T rich repeats associated with constitutive heterochromatin (Celniker et al., 2002; Donnelly and Kiefer, 1986; Hoskins et al., 2015), and this is associated with enhanced Rhi binding to these repeats (Figures 1K and S1G). The *sim-* Cuff ortholog thus leads to reduced RDC binding at clusters, and ectopic binding to constitutive heterochromatin.

### *D. simulans* Cuff traps a cluster transcription complex

To identify protein interactions that are altered by amino acid substitutions in the *D. simulans* ortholog, we expressed GFP tagged *sim-*Cuff and *mel-*Cuff in wild type *D. melanogaster* ovaries, immuno-affinity purified the fusion proteins using GFP-Trap beads, and identified differentially bound proteins by mass spectrometry. To quantify binding, we calculated the ratio of iBAQ values for bound proteins relative to the GFP tag (Figures 3A and 3B). Under our precipitation conditions, which do not use cross-linkers, known piRNA pathway proteins did not co-precipitate with *mel-*Cuff (Figure 3A). However, Cuff colocalizes with Del, and interacts with Del in yeast two-hybrid assays (Mohn et al., 2014). Together, these observations suggest that Cuff directly interacts with Del, but binding is relatively weak and does not survive our immunoprecipitation protocol. In striking contrast, Del was the second most abundant protein, following Cuff itself, in precipitates of *sim-*Cuff (Figures 3A and 3B). In addition, TRF2, which functions with Del and Rhino to promote non-canonical cluster transcription, was the fourth most abundant co-precipitating protein. Substitutions in the *sim-*Cuff protein thus appear to stabilize binding to the *D. melanogaster* Del-TRF2 complex. Rhino did not co-precipitate with *sim-*Cuff or *mel-*Cuff, but Del and Rhino consistently co-precipitate (Figures 3C and 3D). These findings suggest that Del forms separate stable complexes with Cuff and Rhi.

**Figure 3:**
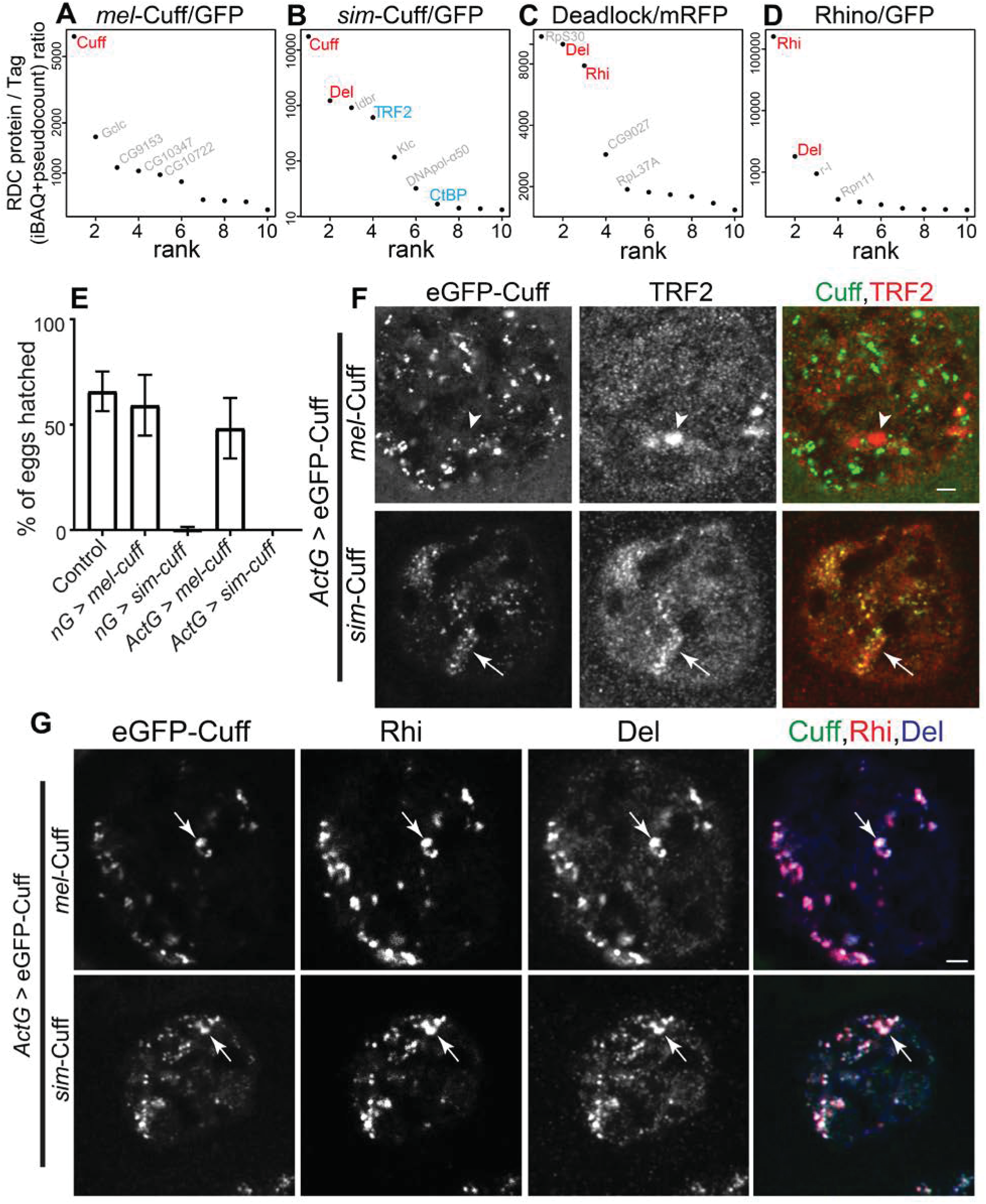
*D. simulans* Cuff traps transcription factors and acts as a dominant negative. (A-D) Mass spectrometric analysis of *mel*-Cuff (A), *sim*-Cuff (B), Del (C) and Rhi (D) binding proteins. Graphs show ratios of iBAQ value of a bound protein in a RDC protein IP vs. tag control IP ranked by ratio values. RDC components are shown in red, TRF2 and CtBP in blue. (E) Bar graphs showing percentages of hatched eggs produced by control (*w^1^*; *Sp*/CyO), flies over-expressing either *mel*-*cuff* or *sim*-*cuff* by either *nanos*-Gal4 (nG) or *Act5C*-Gal4 (*Act*-Gal4) drivers. Error bars show standard deviation of three biological replicates, with a minimum of 200 embryos scored per replicate, except for *nanos*-Gal4 driven *sim-cuff* where average of 50 eggs were scored. (F) Localization of over-expressed GFP tagged Cuff with respect to TRF2 in the germline nuclei of *Act*-Gal4 driven *mel*-Cuff or *sim*-Cuff. Color assignments for merged images shown on top. Arrowheads and arrows denote locations of TRF2 foci. Scale bar, 2 µm. (G) Localization of over-expressed GFP tagged Cuff with respect to Rhi and Del in the germline nuclei of *Act*-Gal4 driven *mel*-Cuff or *sim*-Cuff. Color assignments for merged images shown on top. Arrows denote locations of RDC complex foci. Scale bar, 2 µm.

We speculated that enhanced binding to the Del-TRF2 complex could produce a dominant negative protein, and tested this possibility by over-expressing *sim-*Cuff in wild type females, and assaying fertility, piRNA production, gene and transposon expression. Relatively modest 2.6-fold over-expression of *sim-*Cuff or the *mel-*Cuff control, using the germline specific *rhi* promoter, did not alter fertility (Figure 1B). However, 45-fold over-expression of *sim-*Cuff, using the *UASp* promoter and germline specific *nanos*-Gal4 driver, induced maternal-effect lethality and embryonic dorsal appendage defects, which are characteristic of piRNA pathway mutations (Figure 3E and S4A). Over-expression of *mel-*Cuff, by contrast, did not compromise hatch rate or embryo patterning (Figures 3E and S4A). The somatic follicle cells that surround the developing *Drosophila* oocyte express piRNAs, which are produced through a Cuff-independent mechanism. Mutations that disrupt this somatic piRNA pathway arrest oogenesis and lead to production of rudimentary ovaries (Lin and Spradling, 1997). However, over-expression of *sim-*Cuff in the germline and soma, using an *Act5C-*Gal4 driver, did not reduce ovary size or alter embryo production (not shown), relative to germline-specific *sim-*Cuff over-expression. The *sim-*Cuff protein thus appears to disrupt a germline-specific function.

Cuff is required for transposon silencing and piRNA expression, and RNA-seq demonstrates that *sim-*Cuff over-expression disrupts transposon silencing (Figure S4C). However, small RNA sequencing revealed only a modest reduction in transposon and cluster mapping piRNAs (Figure S4D). Over-expression of *sim-cuff* in wild type, like rescue of *cuff* mutants with *sim-cuff*, thus partially uncouples transposon silencing from piRNA biogenesis (Figures 1D, 1H, S4C and S4D). To gain insight into the molecular basis for this unusual combination of phenotypes, we immuno-precipitated over-expressed *sim-*Cuff and *mel-*Cuff, and identified associated proteins by mass-spectrometry. As observed with *rhino* promoter driven *sim-*Cuff, TRF2 co-precipitated with the over-expressed protein (Figure S3A). In addition, C-terminal Binding Protein (CtBP) consistently showed enhanced binding to *sim-*Cuff, although this protein was also detected, but at lower levels, with *mel-*Cuff. CtBP is a conserved transcriptional co-repressor, initially identified as an Adenovirus E1A binding protein, and subsequently implicated in cancer and control of developmentally regulated genes (Boyd et al., 1993; Schaeper et al., 1995; Stankiewicz et al., 2014). We speculated that the dominant effects of *sim*-Cuff may result from stable binding to TRF2, which promotes non-canonical piRNA cluster transcription (Andersen et al., 2017), and to this conserved repressor of canonical transcription.

To determine if stable binding to *sim-*Cuff alters the *in situ* distribution of these transcription factors, we immuno-localized TRF2 and CtBP in *cuff* mutants expressing low levels of *mel-*Cuff or *sim-*Cuff, and in wild type ovaries over-expressing *mel-*Cuff or *sim-*Cuff. In wild type and *cuff* mutants expressing *mel-*Cuff, TRF2 localized to a few large nuclear domains, which did not overlap with RDC foci (Figures S3B and S3C). These large domains may represent histone repeats, which are regulated by TRF2 (Isogai et al., 2007). In *cuff* mutants expressing *sim-*Cuff, by contrast, TRF2 was displaced from these large foci (Figure S3C), and upon over-expression, TRF2 co-localized with *sim-*Cuff (Figure 3F). Available primary antibodies did not allow direct co-localization of TRF2, *sim-*Cuff, Del and Rhi, but in this genetic background *sim-*Cuff, Del and Rhi co-localize (Figure 3G). Over-expression of *sim-*Cuff thus drives TRF2 into nuclear foci with the RDC. CtBP, by contrast, accumulates in the nucleus, but does not localize to foci in any of these backgrounds (data not shown), suggesting that it forms a distinct complex with *sim-*Cuff.

### CtBP inhibits canonical transcription of piRNA clusters and transposons

TRF2 functions with Del to drive non-canonical cluster transcription (Andersen et al., 2017), but the role of CtBP in the piRNA pathway has not been previously described. *CtBP* null mutants are lethal (Poortinga et al., 1998), so we used RNAi to knock-down *CtBP* in the germline. To confirm specificity, we used 3 different *CtBP* knock-down lines, and a *white* knock-down (*w*-kd) control. The VDRC KK107313 line showed the strongest knock-down efficiency (Figure S5A), and the data obtained using this line are shown in the figures. The vast majority (89.5%) of eggs produced by control *w*-kd females hatch, but *CtBP* knock down reduced the hatch rate to 0.5% (Figure S5B). Strand-specific RNA sequencing showed that this was associated with significant over-expression of 13 transposon families, but only modest changes in gene expression (Figure 4A and S5C). This pattern is typical of piRNA pathway mutations. However, small RNA-seq showed that *CtBP* knock down produced only subtle reductions in cluster and transposon mapping piRNAs (Figures 4C and 4D). piRNA precursor transcripts also showed modest reductions (Figure 4B). *CtBP* knock-down thus mimics *sim-*Cuff over-expression, partially uncoupling transposon silencing from piRNA biogenesis. These findings suggest that *sim-*Cuff binding may inhibit CtBP, triggering dominant sterility.

**Figure 4:**
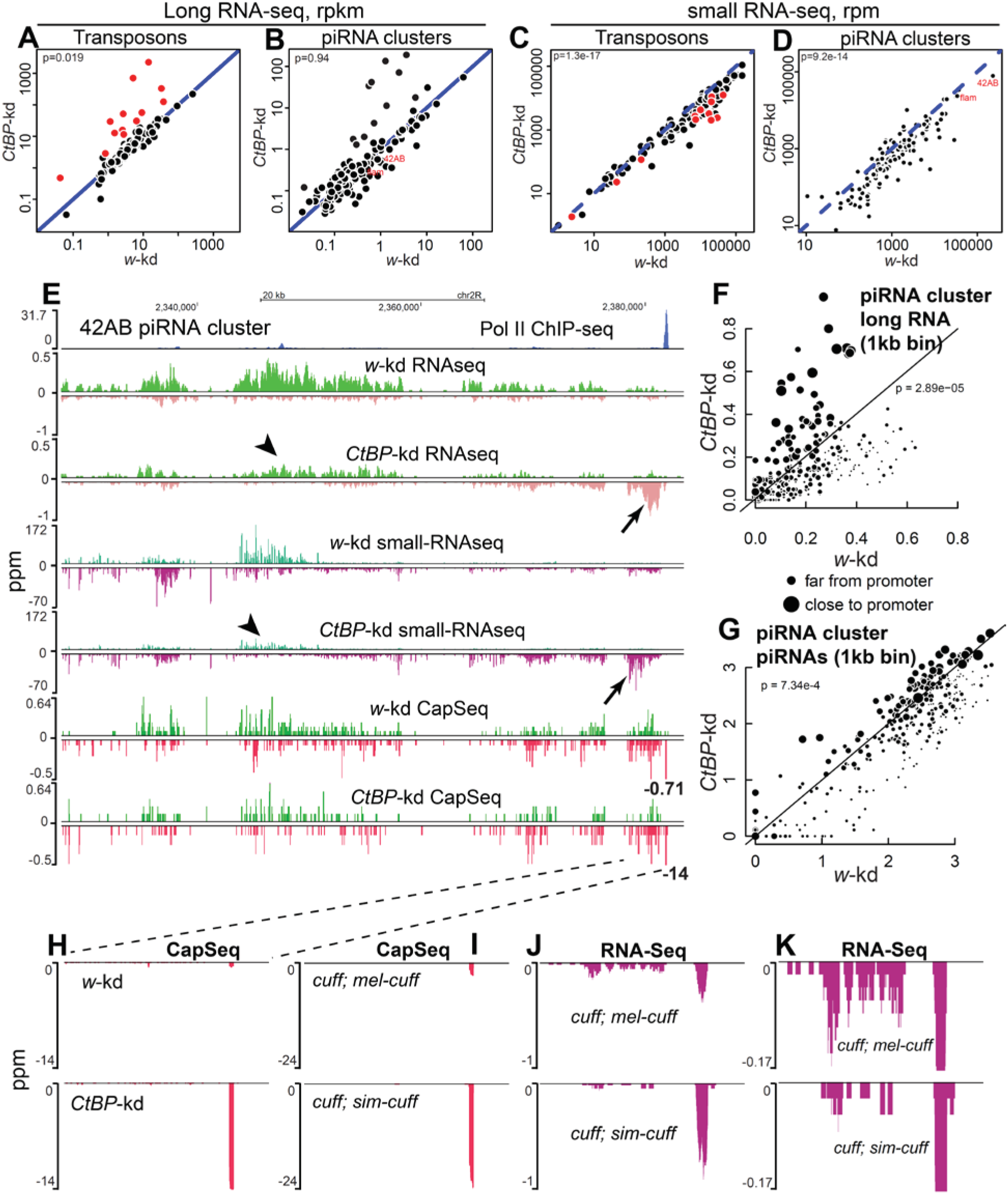
CtBP suppresses canonical transcription at piRNA clusters. (A, B) Scatterplots showing comparisons of RNA-seq signal for transposons (A) and piRNA clusters (B) in *CtBP-*kd vs. *w-*kd ovaries. TEs with more than 3 fold over-expression in *CtBP*-kd vs. *w-*kd as shown in red. (C, D) Scatterplots showing comparisons of small RNA-seq signal for transposons (C) and piRNA clusters (D) in *CtBP-*kd vs. *w-*kd ovaries. Red points denote small RNA expression of TEs, which are over-expressed in *CtBP-*kd as shown in (A). Each point on the scatterplots shows rpkm or rpm value for a transposons family or a piRNA cluster. Diagonal represents x=y. p value for differences obtained by Wilcoxon test. (E) Genome browser view of RNA-seq (top), small RNA-seq (middle) and CapSeq (bottom) profiles at 42AB piRNA cluster from *w-*kd and *CtBP-*kd ovaries. Pol II ChIP-seq peak in *nanos*-Gal4 driven *mel*-Cuff ovaries marks the cluster promoter (blue). Arrows and arrowheads show the increase in canonical transcripts and decrease in non-canonical transcripts respectively after *CtBP-*kd. CapSeq profiles are saturated at promoters. The peak heights of CapSeq promoters are denoted by numbers next to the peaks. (F, G) Scatterplots showing comparisons of rpm values for 1kb bins of piRNA clusters, which have RNA Pol II and TBP promoter peaks, for RNA-seq (F) and small RNA-seq (G) in *CtBP-*kd vs. *w-*kd. The bins close to promoters are shown by big circles and ones farther away by small circles. p value for differences obtained by Wilcoxon test. (H-K) Genome browser views of CapSeq or RNA-seq signals at 42AB promoter for *CtBP-*kd vs. *w-*kd (H) and *cuff* mutants expressing either *mel-cuff* or *sim-cuff* (I-K). (J) and (K) show RNA-seq profiles at different scales.

Most germline piRNA clusters are transcribed from internal non-canonical sites, but the right end of the 42AB cluster and both ends of the 38C cluster are transcribed from canonical promoters, which are marked by prominent RNA Pol II and TATA binding protein (TBP) ChIP-seq peaks (Figures 4E, S5E, 6A and 6B). *CtBP* knock-down produced relatively modest changes in total cluster transcript and piRNA levels, but long RNA and piRNA distributions near the promoters flanking the 42AB and 38C germline clusters were altered (Figures 4E and S5E). Close to the right end of 42AB, *CtBP* knock down produced a significant increase in minus strand long RNAs and piRNAs, and a corresponding decrease in long RNAs and piRNAs from both strands in regions further downstream. A similar pattern was present at both ends of 38C, where plus-strand long RNAs and piRNAs increased at the left flank, while minus strand long RNAs and piRNAs increase at the right flank (Figure S5E). To quantify these observations, we divided the 42AB and 38C clusters into 1kb bins and generated a scatterplot comparing expression in each bin in *w-*kd and *CtBP-*kd, with point size decreasing with increasing distance from the flanking promoters (Figures 4F and 4G). For both long RNAs and piRNAs, *CtBP-*kd increased expression in bins close to promoters (large points), and decreased expression in bins away from promoters (small points). By contrast, the 80F cluster lacks flanking canonical promoters, and *CtBP*-kd did not change long RNA or small RNA expression at this cluster (Figure S5F).

To directly investigate the impact of CtBP on transcription initiation, we used CapSeq to quantify capped transcripts. On *CtBP-*kd, we observed a pronounced increase in capped transcripts associated with promoters flanking 42AB and 38C clusters, and a similar increase in *cuff* mutant ovaries expressing *sim-*Cuff (Figures 4E, 4H and 4I). *CtBP* knock-down or expression of the *sim-*Cuff protein thus appear to activate canonical promoters flanking 42AB and 38C, which is associated with reduced non-canonical transcription from downstream sequences.

Heterochromatic clusters are the major source of germline piRNAs in *Drosophila* ovaries, but a subset of isolated euchromatic transposons function as “mini piRNA clusters” and are bound by Rhino and produce sense and anti-sense piRNAs (Figures 5A and 5B). Because transposons are repeated, internal sequences cannot be mapped to specific insertions. However, Rhino spreads into flanking unique sequence from the source insertions, and read through non-canonical transcription leads to piRNA production from these unique regions, producing a characteristic “butterfly” piRNA profile. To identify these loci, we used paired end genome sequencing to map all euchromatic transposon insertions, and then identified insertions with flanking Rhino ChIP-seq peaks and divergently expressed piRNAs. Figures 5A and 5B show examples of *Diver* and *Blood* insertions that function as mini-clusters in the *w*-kd line. In both cases, *CtBP*-kd reduced Rhino binding, and triggered a near collapse of flaking piRNA expression. The scatter plots in Figures 5C and 5D summarize data for all of the new piRNA producing insertions we identified with genomic DNA sequencing, and show that this pattern extends across the genome. In addition, CapSeq shows that the loss of Rhino binding and piRNA production is associated with significant increases in canonical transcription from promoters within the LTRs of the inserted elements (Figure 5E). By contrast, transcription initiation from within the transposons, which appears to reflect non-canonical transcription, is reduced for both sense and anti-sense strands (Figures 5F and 5G). CtBP thus suppresses canonical transcription from promoters linked to clusters and euchromatic transposon insertions. In both contexts, increased canonical transcription is associated with reductions in both non-canonical transcription and piRNA production.

**Figure 5:**
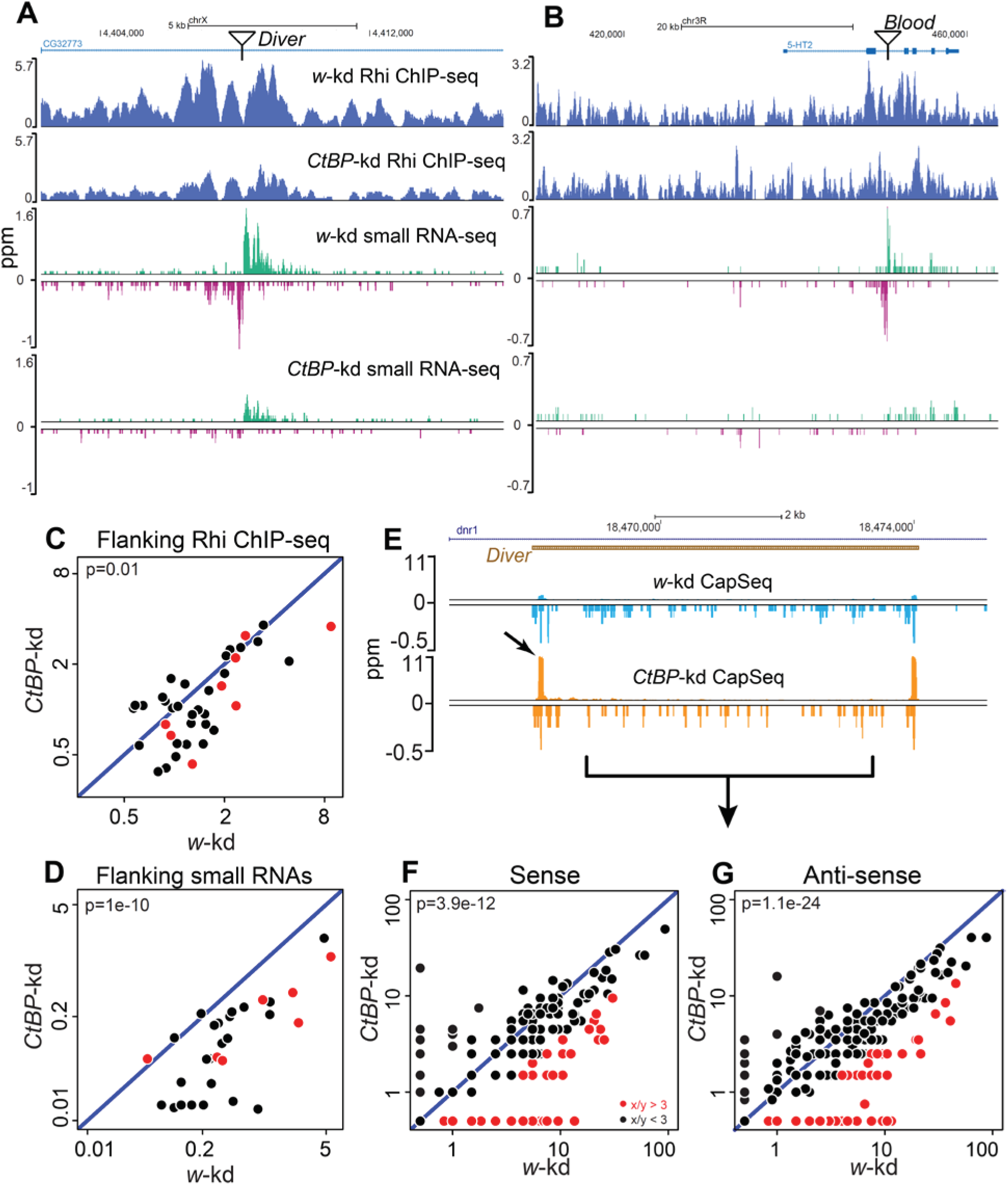
CtBP suppresses canonical transcription of dispersed transposon insertions. (A, B) Genome browser views of Rhino ChIP-seq and small RNA-seq profiles flanking dispersed transposons, *Diver* (A) and *Blood* (B), in *CtBP-*kd and *w-*kd. The transposon insertion is shown at the top. (C, D) Scatterplots showing comparisons of rpm values of Rhi ChIP-seq and small RNAs, 0.5kb upstream and downstream of new transposons in *CtBP-*kd vs. *w-*kd. The transposons insertions were identified by genomic sequencing with TEMP (Zhuang et al., 2014) and the graphs show the values for new TEs (not present in the reference genome), which have both flanking piRNAs and Rhi signal. Red points denote expression of TEs over-expressed in *CtBP-*kd, as shown in Figure 4A. p value for differences obtained by Wilcoxon test. (E) Genome browser view of CapSeq signal at *Diver* insertion in *CtBP-*kd vs. *w-*kd. Arrow shows increased CapSeq signal at *Diver* 5’ end in *CtBP-*kd. (F, G) Scatterplots showing comparisons of CapSeq signal for 1kb bins mapping to transposons present outside clusters, (except for the bins at 5’ and 3’ ends, to remove canonical transcription peaks) for *CtBP-*kd vs. *w-*kd. (F) shows sense strand and (G) shows anti-sense strand initiation. Points in red show x/y>3. p value for differences obtained by Wilcoxon test.

### Cuff associates with canonical and non-canonical transcription sites

These data, with previous studies, link Cuff to factors that regulate canonical and non-canonical transcriptions of piRNA source loci. Further supporting this link, our ChIP-seq analysis shows that endogenous Cuff localizes with Pol II and TATA binding protein (TBP) at canonical promoters flanking major germline clusters, and confirms earlier data showing that Cuff co-localizes with Rhino and Del at sites of non-canonical transcription in the body of piRNA clusters (Figures 6A, with 6B being the zoomed-in view of the canonical transcription start in 6A). Cuff, Rhi and Del are co-dependent for cluster binding (Chen et al., 2016; Mohn et al., 2014). Consistent with these studies, long RNA and CapSeq indicate that *cuff* mutations significantly reduce transcription from both strands of internal cluster sequences (Figure 6A), and ChIP-seq indicate that this correlates with reduced Rhi binding to 42AB and other germline piRNA clusters (Figures S6A and S6B). By contrast, *cuff* mutants did not reduce CapSeq signal associated with the canonical promoters flanking the 42AB (Figure 6B) or 38C clusters (Figure S6C). However, the transcripts from these canonical promoters are terminated shortly after initiation (Figure 6B), and tethering *Cuff* to the 3’ end of a reporter transcript enhances read through transcription (Chen et al., 2016). These findings suggest that endogenous Cuff suppresses termination of transcription from canonical promoters flanking the major germline clusters, but does not directly regulate transcription initiation from these promoters. By contrast, rescue of *cuff* mutants with the *sim-*Cuff ortholog leads to a 7.7 and 2.3 fold expression of capped transcripts from the 42AB and 38C promoters respectively (Figures 4I and S6C). We speculate that this increase is due *to sim*-Cuff binding to CtBP, leading to partial inhibition.

**Figure 6:**
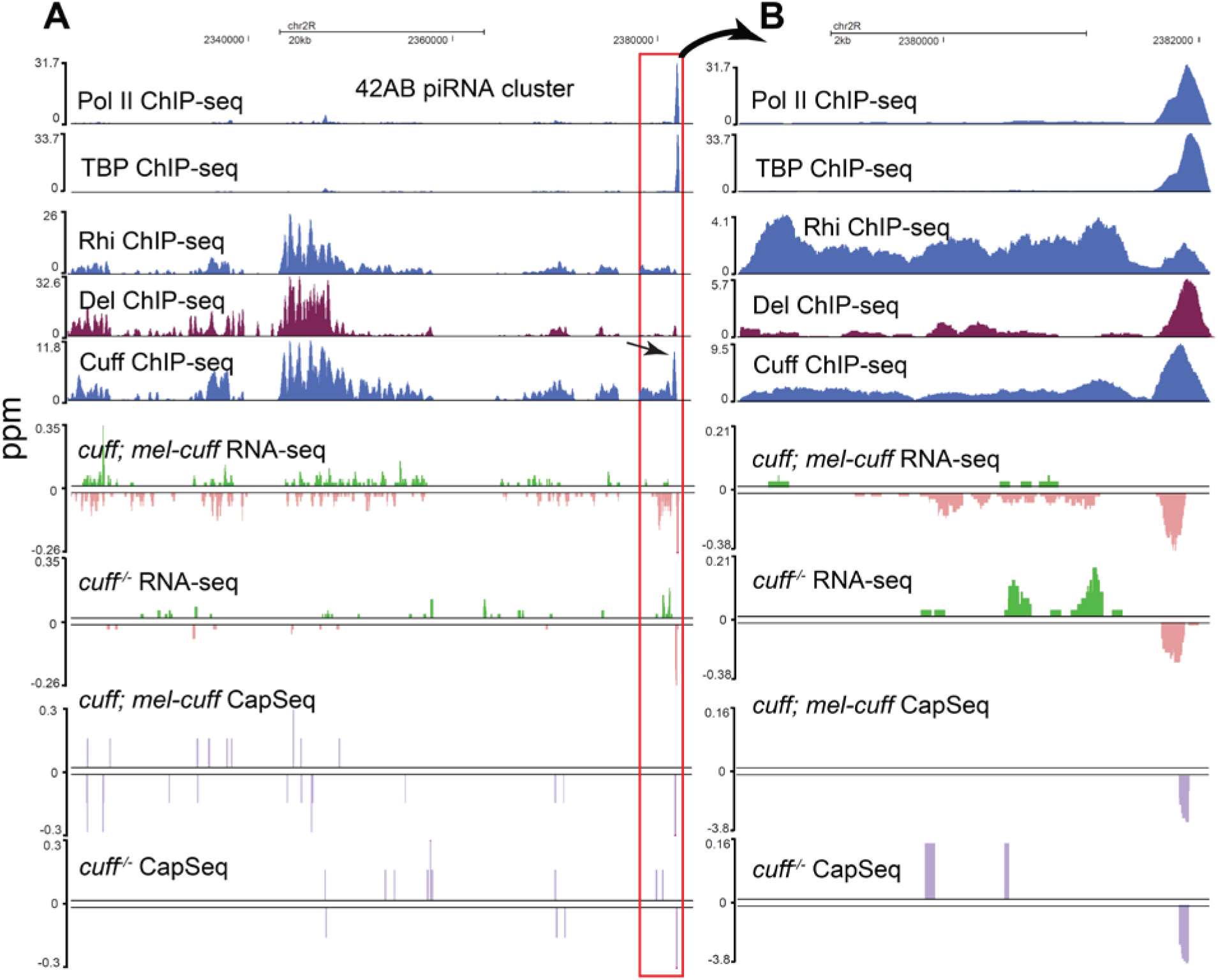
Role of Cuff in piRNA cluster transcription. (A, B) Genome browser views at 42AB cluster. (A) Right side of the 42AB cluster, proximal to the flanking canonical promoter, showing Pol II, TBP (TATA-binding protein), Rhi, Del and Cuff ChIP-seq, RNA-seq and CapSeq signals. Rhi, Del and Cuff localize throughout the clusters, while Cuff and Del also show peaks that correspond to the flanking canonical promoter, marked by Pol II (arrow). (B) Zoomed in view of the promoter region for all the tracks in (A). All the ChIP-seq tracks are auto-scaled. RNA-seq and CapSeq profiles shown in *cuff* mutants and *cuff* mutants expressing *mel-cuff*.

## DISCUSSION

Adaptive evolution is characteristic of genes engaged in a genetic conflict, and can result from positive selection of pathogen mutations that reduce binding of host defense proteins and allow proliferation, followed by selection of host mutations that restore binding and pathogen control (Daugherty and Malik, 2012). The resulting arms race drives rapid co-evolution of host-pathogen gene pairs. However, pathogens can produce “mimics” of host proteins, which compete for productive binding between host proteins in the defense machinery (Elde and Malik, 2009). Adaptation to mimics can remodel host protein-protein interactions, producing orthologs that are binding site mutations when moved across species, providing a novel approach to structure-function analysis (Parhad and Theurkauf, 2019). We previously used this approach to define an interaction between the Rhi Shadow domain and Del that prevents ectopic piRNA cluster chromatin assembly (Parhad et al., 2017; Yu et al., 2018). Here we show that adaptive evolution of the third RDC component, *cuff*, targets interactions with conserved proteins that regulate piRNA cluster transcription and piRNA biogenesis.

### *sim-*Cuff captures piRNA precursor transcription factors

Transposon silencing piRNAs are derived from heterochromatic clusters and a subset of euchromatic transposon insertions, and Cuff co-localizes with Rhi and Del at these piRNA source loci (Mohn et al., 2014). Rhi binds to H3K9me3 marks and recruits Del, which interacts with Moonshiner and TRF2 to initiate non-canonical transcription from both genomic strands (Andersen et al., 2017; Le Thomas et al., 2014; Mohn et al., 2014; Yu et al., 2015). By contrast, the role of Cuff is less well understood. However, the *cuff* gene is evolving rapidly, suggesting that cross-species studies may provide novel functional insights. We show that *D. simulans cuff* ortholog does not complement *D. melanogaster cuff* mutations (Figure 1), and that *sim-*Cuff over-expression triggers dominant sterility and transposon over-expression (Figures 3 and S4). Significantly, this dominant activity is associated with assembly of stable complexes with Del and TRF2, which drive non-canonical transcription, and to CtBP, which is a conserved co-repressor of canonical transcription.

Dominant mutations can produce new interactions or functions (neomorphic mutations), triggering assembly of complexes that are not formed by wild type proteins (Jeffery, 2011). However, our data indicate that the *sim-*Cuff ortholog, when placed in a *D. melanogaster* background, stabilizes transient interactions with Del, TRF2 and CtBP, which function with Cuff to regulate piRNA precursor transcription. In *D. melanogaster*, Cuff and Del do not co-precipitate. However, these proteins co-localize, interact in two-hybrid assays, and are co-dependent for association with piRNA clusters (Mohn et al., 2014). Del, in turn, co-precipitates with TRF2 and Moonshiner, all three proteins are required for non-canonical cluster transcription (Andersen et al., 2017). Significantly, we show that Cuff is necessary for non-canonical transcription, and that CtBP suppresses canonical transcription of transposons and promoters flanking the major germline clusters. Adaptive evolution has therefore altered Cuff binding to a suite of proteins that control canonical and non-canonical piRNA precursor transcription.

Competition between canonical and non-canonical transcription appears have a central role in controlling piRNA biogenesis (Andersen et al., 2017; Chang et al., 2019), and is particularly critical to piRNA production from euchromatic transposon insertions. The majority of germline clusters do not have flaking canonical promoters, and CtBP knock down does not significantly alter piRNA production from these loci. However, canonical promoters flank the right side of the 42AB cluster and both ends of the 38C clusters, and at these clusters CtBP knock down and promoter activation reduces non-canonical transcription and piRNA expression from downstream regions. Perhaps more significantly, CtBP knock down leads to a near collapse of piRNA production and Rhi binding at euchromatic transposon insertions that function as “mini-clusters” (Figure 5). We cannot directly assay non-canonical transcription at specific transposon insertions, as internal sequences are repeated, but the loss of piRNAs from flanking unique sequences implies that non-canonical transcription is blocked. Deletion of the promoters flanking 42AB and 38C leads to spreading of piRNA production into flanking domains (Andersen et al., 2017). The data presented here, with these earlier observations, indicate that canonical transcription directly or indirectly represses non-canonical transcription and piRNA production.

### A piRNA precursor transcription network

These observations lead us to propose that Cuff functions in a network that regulates the balance of canonical and non-canonical transcription of piRNA source loci (Figure 7). At large heterochromatic clusters composed of nested transposon arrays, and the subset of recent euchromatic transposon insertions that produce piRNAs, Cuff functions with Del and TRF2 to promote non-canonical transcription and piRNA production from both genomic strands. As mutations in *cuff* nearly eliminate non-canonical transcription, it led us to propose that Cuff is limiting for assembly of a transient complex containing Cuff, Del and TRF2, which binds Rhi and drives non-canonical transcription initiation. Active transposon insertions and the two major clusters at 42AB and 38C are also transcribed from canonical promoters, which are suppressed by CtBP. Transcription from these canonical promoters, in turn, suppresses downstream non-canonical transcription and piRNA biogenesis. This likely reflects transcriptional interference, resulting from displacement of the non-canonical transcription machinery by elongating RNA Polymerase II. Competing canonical and non-canonical transcription appears to be coordinated through Cuff-CtBP and Del-TRF2-Cuff complexes, which is stabilized by the *sim-*Cuff ortholog. Stable binding to *sim-*Cuff leads to sterility, enhanced canonical transcription, and reduced piRNA production. We therefore propose that this complex sequesters CtBP and stabilizes Del-TRF2 complex, thereby activating both canonical and non-canonical transcription. Under normal conditions, these complexes are unstable, freeing CtBP to suppress canonical promoters and Del-TRF2 to start new rounds of non-canonical transcription. However, environmental stress, which can lead to transposon activation (Maze et al., 2011; Miousse et al., 2015; Natt and Thorsell, 2016), may promote assembly of these complexes. CtBP is an NADH/NAD binding protein (Fjeld et al., 2003; Jack et al., 2011), raising the intriguing possibility that this network couples piRNA biogenesis and transposon silencing to metabolic state.

**Figure 7:**
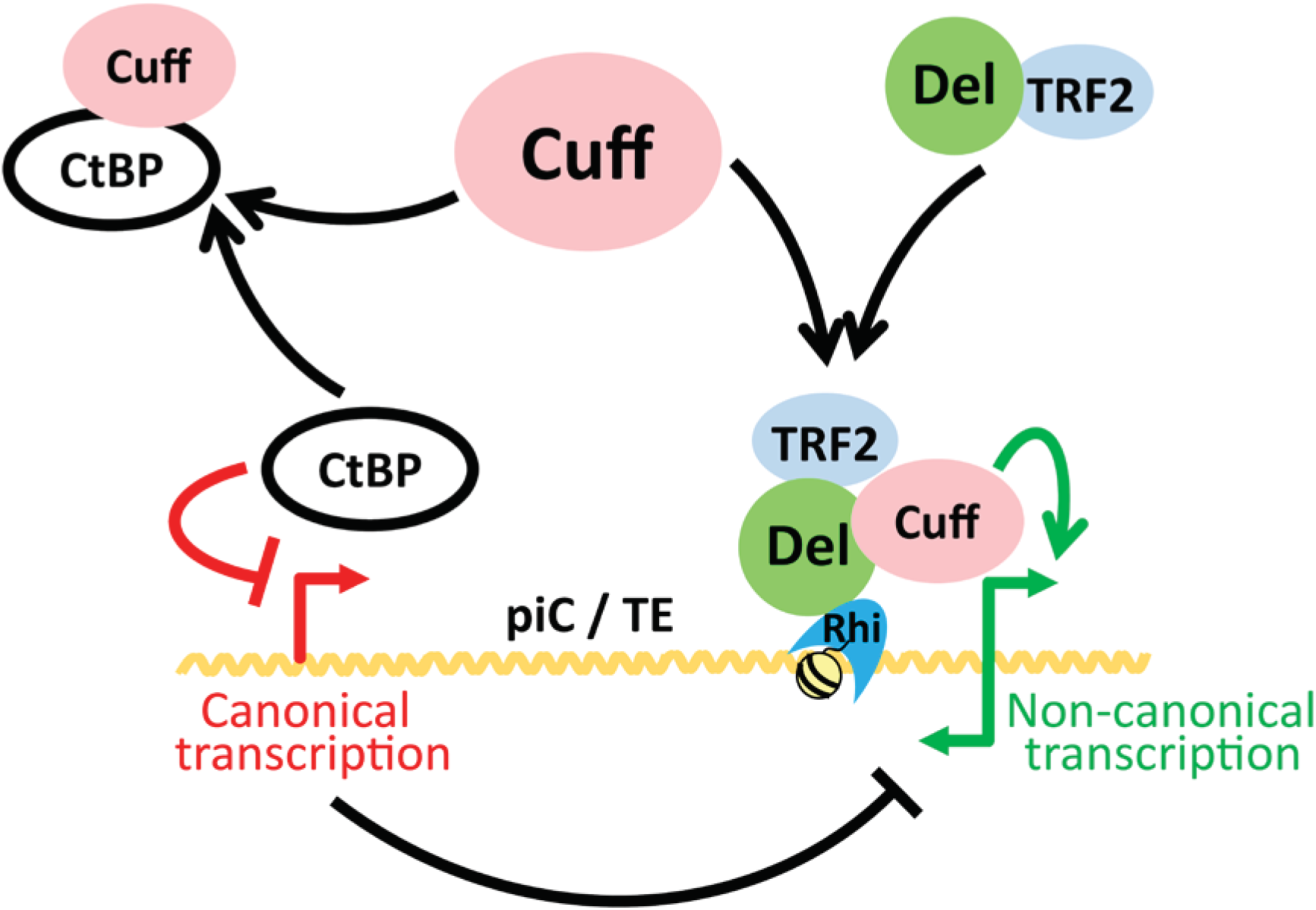
Model for a transcriptional network balancing canonical and non-canonical piRNA precursor transcription. piRNAs are generated from both piRNA clusters and dispersed transposon insertions, which act as “mini-clusters”. At both locations, Rhino binds to H3K9me3 histone marks and recruits Del-TRF2-Cuff complex to initiate non-canonical transcription from both strands. Non-canonical transcription (green lines) is inhibited by canonical transcription (red lines), and CtBP represses canonical transcription, regulating non-canonical transcription and piRNA production.

All three RDC genes are rapidly evolving and have accumulated substitutions that alter interactions with binding partners, which do not function across species. Rapid evolution of host protein-protein interactions, as opposed to host-pathogen gene pairs, is consistent with adaption to a molecular mimic (Elde and Malik, 2009). Transposons are the most likely source for mimics targeting piRNA biogenesis, but numerous piRNA pathway genes are evolving rapidly, and the genetic repertoire of simple transposons is limited (Parhad and Theurkauf, 2019). It seems unlikely that all of the rapidly-evolving piRNA pathway genes are targeted by different mimics. However, these genes function within networks, and we speculate that a single mimic targeting one interaction could drive compensatory changes in biochemically coupled proteins. For example, Cuff binding to Del-TRF2 could produce a Del-TRF2-Cuff complex that interacts with Rhi, presumably through Del, which drives non-canonical transcription. A mimic competing for Rhi binding to Del would reduce formation of non-canonical transcription complexes on chromatin, which could be restored by mutations in Del that increase affinity and displace the mimic. Alternatively, mutations in Cuff that increase binding to Del-TRF2 would increase the concentration of the Del-TRF2-Cuff complex, driving increased binding to Rhi and non-canonical transcription. In this model, a single mimic could drive “coupled evolution” of multiple interactions, leading to multiple incompatibilities between piRNA pathway proteins from closely related species. This could explain why sterile hybrids between *D. simulans* and *D. melanogaster* phenocopy piRNA pathway mutations (Kelleher et al., 2012; Sturtevant, 1919, 1920).

All three RDC components are rapidly evolving and have no clear homologs outside of *Drosophila.* By contrast, TRF2 and CtBP are conserved from flies to humans (Chinnadurai, 2002; Rabenstein et al., 1999), and CtBP is a putative human oncogene (Dcona et al., 2017; Stankiewicz et al., 2014). Rapidly evolving genes with specialized functions are frequently the most accessible to phenotype based forward genetic approaches in model systems, and linking these specialized genes to conserved pathway components can be a challenge. The studies reported here indicate that cross-species studies can link rapidly evolving genes to conserved factors, and bridge the gap between studies on genetically-tractable model organisms and humans.

## AUTHOR CONTRIBUTIONS

S.S.P., W.E.T. and Z.W. conceived the project. S.S.P. performed all the experiments, except *CtBP*-kd and *w*-kd RNA-seq and small RNA-seq performed by G.Z. and N.P.R. respectively. S.S.P. and T.Y. performed bioinformatics analysis. S.S.P. and W.E.T. wrote the paper with help of Z.W. and other authors.

## ACKNOWLEDGEMENTS

We would like to thank the members of Weng and Theurkauf labs and the UMass RNA Biology community for their insightful discussions and critical comments; Trudi Schüpbach for *cuff* stocks; Julius Brennecke for Del antibody; James Kadonaga for TRF2 antibody; Bloomington, VDRC and UCSD *Drosophila* stock centers for fly stocks; John Leszyk and the UMass Proteomics facility for mass spectrometry. This work was supported by National Institute of Child Health and Human Development (R01HD049116 and P01HD078253).

## DECLARATION OF INTERESTS

The authors declare no competing interests.

## KEY RESOURCES TABLE

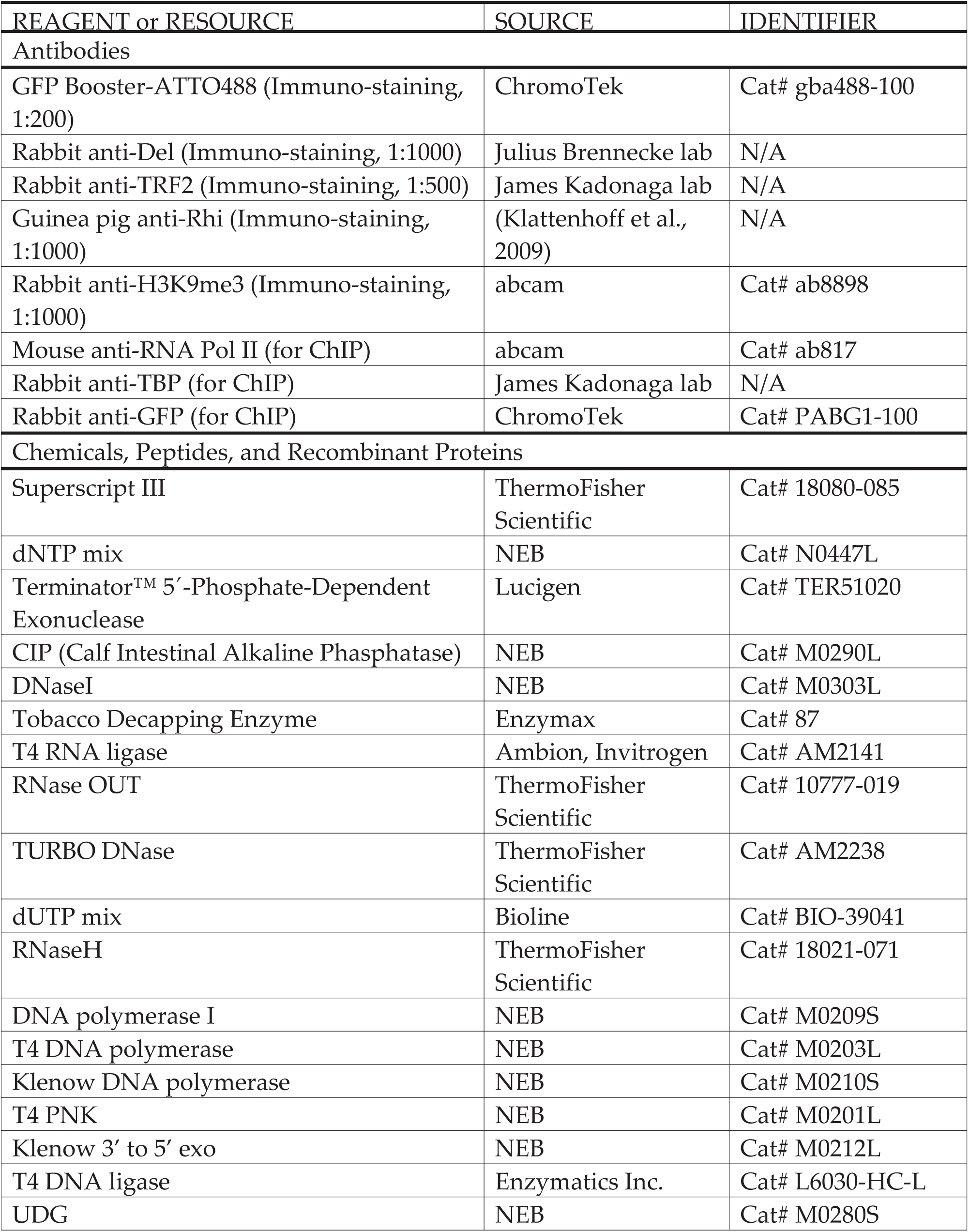
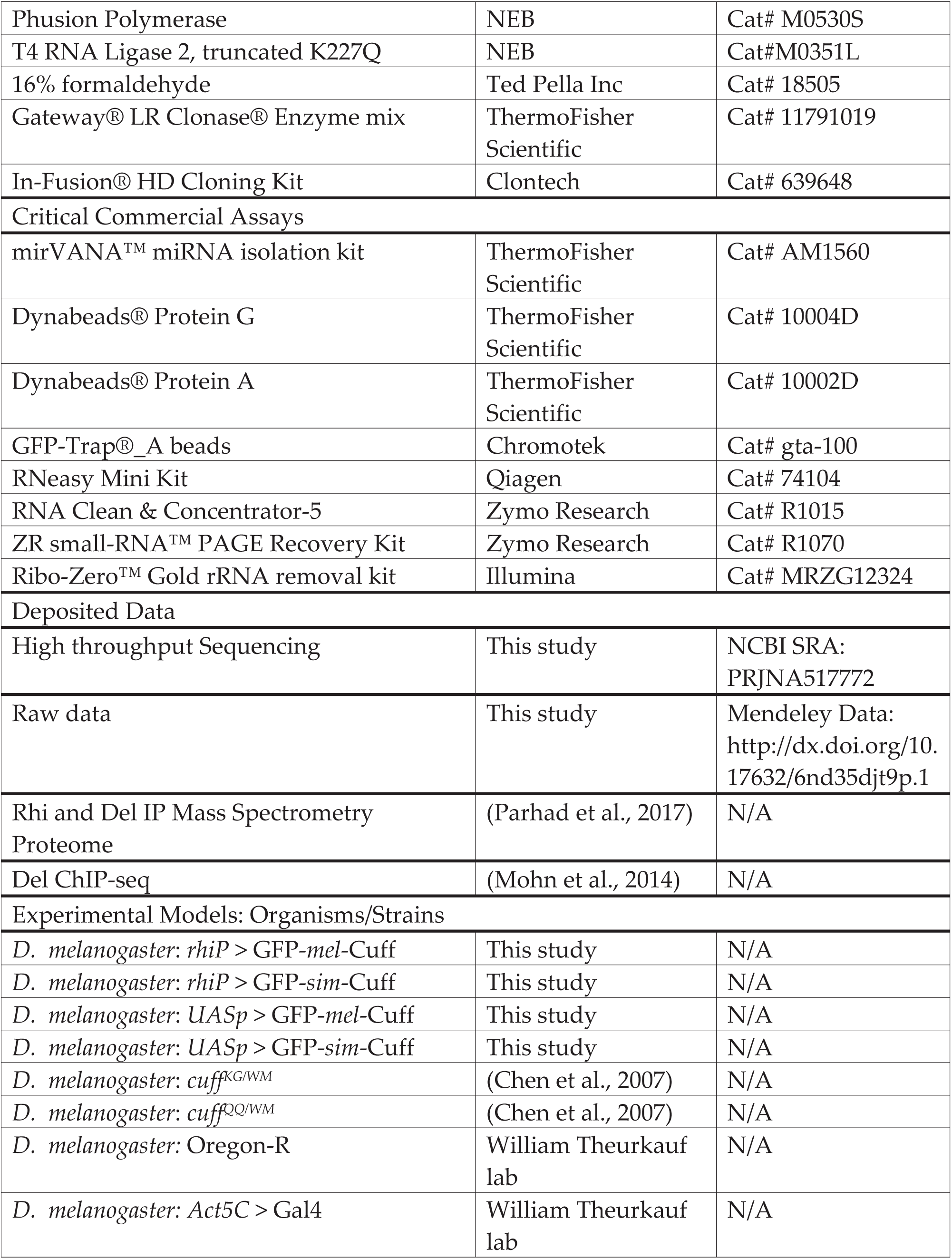
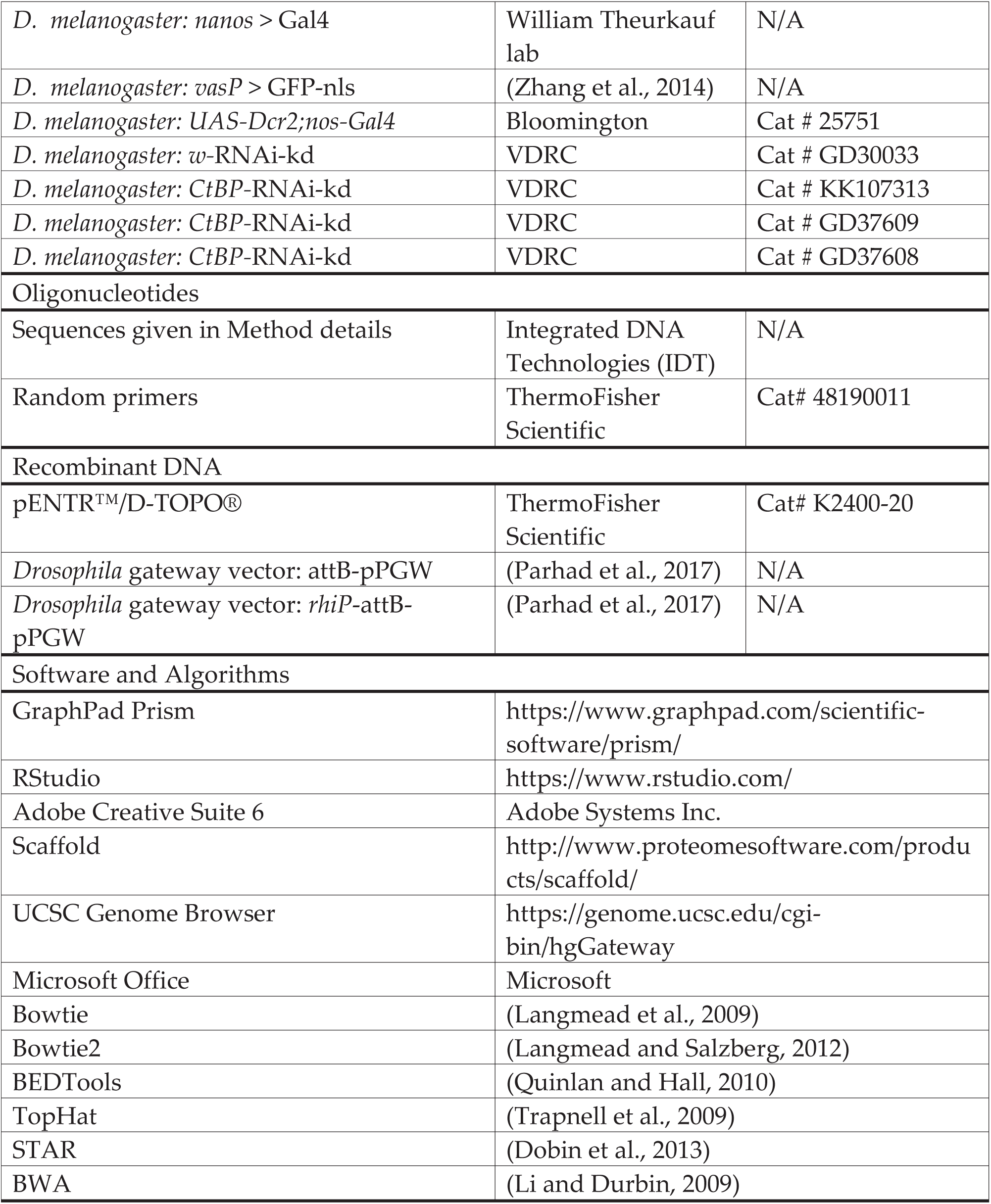

## STAR METHODS

### CONTACT FOR REAGENT AND RESOURCE SHARING

Further information and requests for resources and reagents should be directed to and will be fulfilled by the Lead Contact William Theurkauf.

### EXPERIMENTAL MODEL AND SUBJECT DETAILS

All experiments were performed in 2-4 day old *Drosophila melanogaster* females, except mentioned otherwise. All flies were maintained at 25°C on cornmeal medium. All transgenic lines were generated by φC31 integration at 3L-68A4. *cuff*^*WM25*^ (*cuff*^*WM*^) and *cuff*^*QQ37*^ (*cuff*^*QQ*^) alleles were obtained from Trudi Schüpbach (Princeton University) (Chen et al., 2007). *cuff*^*KG05951*^ (*cuff*^*KG*^) was obtained from Bloomington (Stock # 14462). *Act5C*-Gal4 and *nanos*-Gal4 stocks were used from our lab stocks. RNAi knockdown lines were obtained from VDRC.

### METHOD DETAILS

#### Generation of transgenic flies

*mel-cuff* was cloned from *D. melanogaster* OregonR ovary cDNA and *sim-cuff* from *Drosophila simulans* C167.4 ovary cDNA. The reverse primer for the PCR reaction was used for making cDNA with Superscript III reverse transcriptase (Thermo Fisher Scientific). *mel-cuff* was PCR amplified from cDNA by using forward primer: CAC CAT GAA TTC TAA TTA CAC AAT ATT AAA C and reverse primer: TTA AAC TAT AGA AGA CAT GGT TTG C and cloned into pENTR-D-TOPO vector by directional TOPO cloning kit (Thermo Fisher Scientific). Similarly, *sim-cuff* was PCR amplified from cDNA using forward primer: CAC CAT GAA TTC TAA TTA CAA AAT ATT GAA C and reverse primer: TTA TTG GTA AAC TGT GGA AGA CAT GG and cloned into pENTR-D-TOPO vector. These served as entry vectors for Gateway cloning. The destination vectors *rhiP*-attB-pPGW (for expressing N’ GFP tagged proteins under *rhi* promoter) and attB-pPGW (for expressing N’ GFP tagged proteins under *UASp* promoter) were used as described in (Parhad et al., 2017). The plasmids obtained from LR gateway cloning reaction were sequenced and injected by φC31 integration at chromosomal location 3L-68A4 (Bischof et al., 2007).

#### Fertility assays

2-4 day old flies were maintained on grape juice agar plates for 1 or 2 days. After removing flies, the eggs were counted for fused appendages. The number of hatched eggs were counted after 2 days. The fertility bar graphs indicate mean and standard deviation from 3 biological replicates.

#### RT-qPCR

RNA was isolated from 2-4 day old female ovaries. Reverse transcription done using Superscript III reverse transcriptase with random primers. qPCR was done by Qiagen QantiTect® SYBR® Green PCR mix using Applied Biosystems instrument. Primers sequences for CtBP: forward primer: CAA AAA TCT GAT GAT GCC GAA GCG TTC and reverse primer: AGG ATG GGC ATC TCG ATG GAG CAG TC and Rp49: forward primer: CCG CTT CAA GGG ACA GTA TCT G and reverse primer: ATC TCG CCG CAG TAA ACG C.

#### Immuno-staining

Immuno-staining and image analysis were performed as described in (McKim et al., 2009; Zhang et al., 2012a). In short, 2-4 day old female ovaries were dissected in Robb’s buffer, fixed with 4% formaldehyde, washed, overnight incubated with primary antibody, washed, incubated overnight with secondary antibody with the fluorophore, stained with DAPI for DNA labelling and mounted on slide with mounting medium. To enhance the GFP signal, ChromoTek anti-GFP Booster (Atto-488) antibody was added with secondary antibody. Antibodies used: anti-GFP Booster (ChromoTek) at 1:200, guinea pig anti-Rhi (our lab) at 1:1000, rabbit anti-Del (from Julius Brennecke) at 1:1000, rabbit anti-TRF2 (from James Kadonaga) at 1:500, rabbit anti-H3K9me3 (abcam) at 1:1000.

#### Immuno-precipitation

IP was performed as described in (Parhad et al., 2017). Briefly, 2-4 day old female ovaries were dissected in Robb’s medium, lysed by homogenization and sonication and centrifuged to get input for IP. Lysis and IP buffer composition: HEPES (pH 7.5) 50mM, NaCl 150mM, MgCl2 3.2mM, NP-40 0.5%, PMSF 1mM, Proteinase Inhibitor (Roche) 1X. chromotek GFP-Trap®_A beads were used for GFP IP. The lysate was incubated with beads for 3 hours at 4°C and subsequently washed with lysis buffer 4 times. Finally the beads were suspended in SDS-PAGE lysis buffer. The procedure for mass spectrometry is descried in (Vanderweyde et al., 2016). Briefly, the IP samples were resolved on a 10% SDS-PAGE gel. The gel pieces were trypsin digested to get the peptides, which were analyzed by LC-MS/MS. Rhi and Del IP data was used from (Parhad et al., 2017).

#### Small RNA-seq

Small RNA libraries were prepared as mentioned in (Zhang et al., 2014). In short, total RNA was prepared by mirVANA kit (Ambion). 18-30 nt length small RNAs were size selected by denaturing PAGE gel purification. These were ligated further at 3’ and 5’ ends by adapters. Reverse transcription and then PCR amplification was performed to obtain libraries. Single end sequencing was done by Illumina platform.

#### RNA-seq

RNA-seq libraries were prepared as described in (Fu et al., 2018; Zhang et al., 2018; Zhang et al., 2012b). Briefly, RNA samples were depleted for ribosomal rRNA by Ribo-Zero kit (Illumina) or rRNA digestion by RNaseH (Epicenter), fragmented and reverse transcribed. After dUTP incorporation for strand specificity, end repair, A-tailing, adapter ligation and PCR amplification was done to obtain libraries. Paired end sequencing was done by Illumina platform.

#### CapSeq

This method was performed to sequence 5’ ends of transcripts (Gu et al., 2012). In brief, total RNA was sequentially treated with Terminator™ 5′-Phosphate-Dependent Exonuclease, CIP (Calf Intestinal Alkaline Phasphatase), DNaseI, Tobacco Decapping Enzyme. After adapter ligation at the 5’ end, reverse transcription (with primer: 5’-GCACCCGAGAATTCCANNNNNNNN-3’) and two rounds of PCR were done. The PCR products were gel purified after each PCR step. Final library was sequenced by Illumina platform by single end sequencing.

#### ChIP-seq

ChIP-seq was performed by method described in (Parhad et al., 2017). In short, the ovaries were dissected in 1X Robb’s medium and fixed with 2% formaldehyde and sonicated in Bioruptor for 2 hours. This lysate was centrifuged and the supernatant was used as input for ChIP. The input was precleared with either Dynabeads Protein A or Dynabeads Protein G (Invitrogen) and was added to the Dynabeads conjugated to an antibody and incubated overnight. After washing, the beads were reverse crosslinked, ChIP DNA was purified and libraries were prepared by end repair, A tailing, adapter ligation and PCR amplification. Illumina platform was used for paired end sequencing.

### QUANTIFICATION AND STATISTICAL ANALYSIS

#### Bioinformatics analysis

Small RNA-seq reads were aligned to the *D. melanogaster* genome (dm3) and transposon consensus sequences by bowtie (Langmead et al., 2009), after removing the 3′end linkers. Flybase r5.50 transcriptome annotations were used. The piRNA cluster coordinates were from (Brennecke et al., 2007). Reads mapping to known non-coding RNAs (ncRNAs, such as rRNAs, tRNAs, etc.) and miRNAs were excluded for the quantification of piRNA abundance. The read counting was done using BEDTools (Quinlan and Hall, 2010) and normalized to microRNAs. Multiple mapping locations are considered while counting reads. TopHat (Trapnell et al., 2009) was used to align RNA-seq reads to the genome. rRNA reads were removed prior to the quantification of genes, piRNA clusters, and transposons expression. For ChIP-seq, alignment was done by Bowtie2 (Langmead and Salzberg, 2012) and normalized to total mapped reads. For CapSeq, we used STAR for mapping (Dobin et al., 2013) after removing the RT primer sequence. Total uniquely mapped reads were used as the normalization factor.

#### Mass spectrometry Proteomic Analysis

Proteome Discoverer and Mascot Server were used to process the raw data before display on Scaffold Viewer (Proteome Software, Inc.). The proteins were filtered by criteria: Protein threshold: 90%, Min # peptides: 2, Peptide threshold: 90%. Then iBAQ values (Schwanhausser et al., 2011) were obtained and pseudocount was added. For Cuff IP, *vas* promoter driven GFP-nls was used as a control. Both replicates of *rhi* promoter (*rhiP*) driven Cuff IP mass spectrometry scaffold tables were combined into a single file. To obtain list of proteins binding to Cuff and not GFP control, only proteins below the threshold of 300000 in GFP IP were selected. The proteins which show more than 3 fold enrichment in all the Cuff protein IPs vs. GFP control IP were used to make plots, where the ratios of (iBAQ + psuedocount) values for each identified protein in a Cuff IP vs. GFP IP were plotted against their rank. For sim-Cuff graphs, in addition to the above filters, proteins which show more than 3 fold enrichment for sim-Cuff IP vs. mel-Cuff IP were plotted. The graphs were made using R. Similar filters and thresholds were used for Rhi and Del IP mass spectrometry data from (Parhad et al., 2017).

#### Analysis of RT-qPCR data

Quantification done using ΔCt method. Rp49 served as the loading control.

#### Statistical Analysis

The error bars in the bar graphs show standard deviations from 3 biological replicates.

### DATA AND SOFTWARE AVAILABILITY

Cloned cuff cDNA sequences and Cuff Proteomics data are deposited in Mendeley Data: http://dx.doi.org/10.17632/6nd35djt9p.1. High throughput sequencing data can be accessed from NCBI SRA: PRJNA517772.

---

**Figure S1, related to figure 1:**
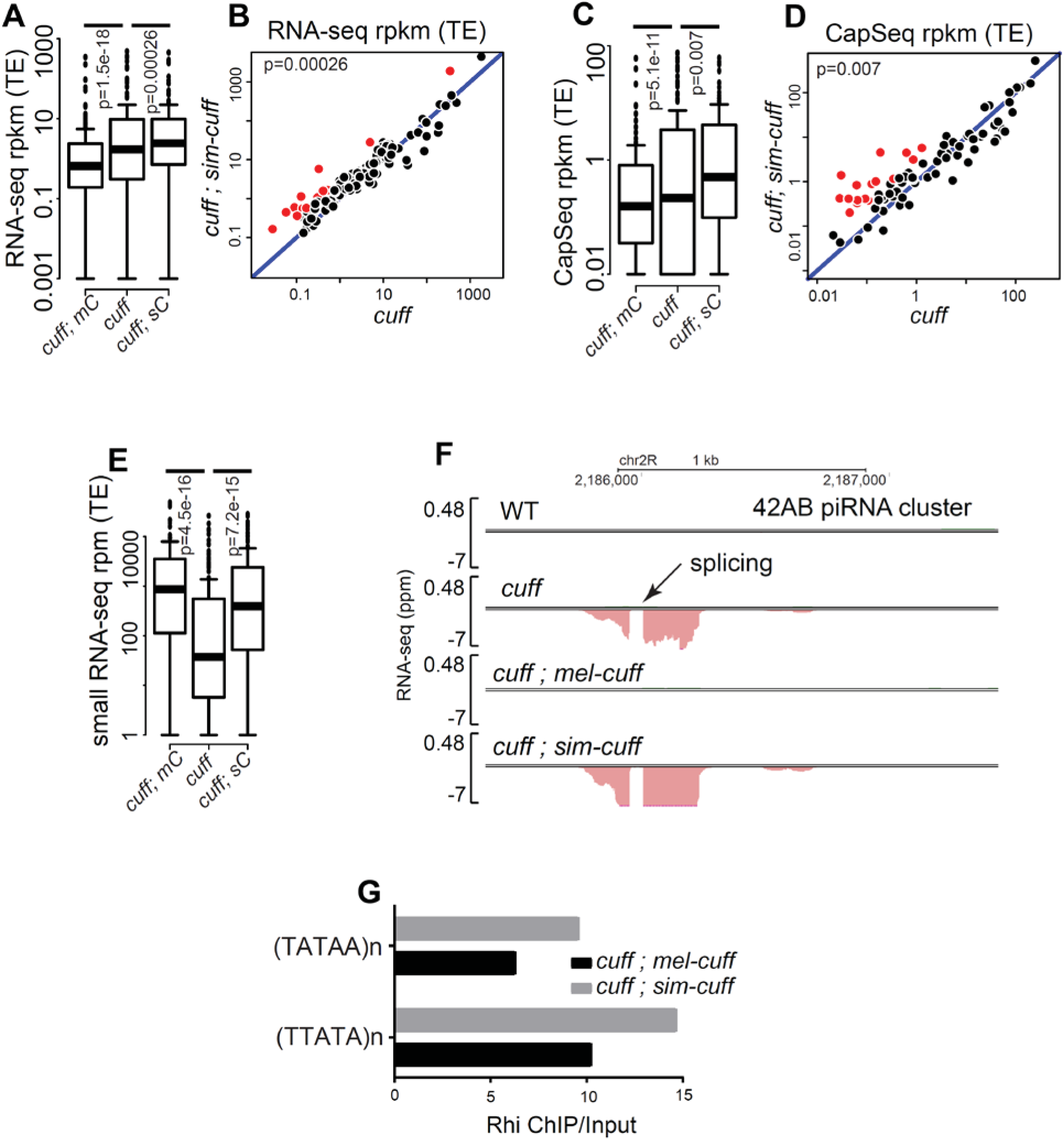
(A-E) Boxplots and scatterplots showing transposon mapping long-RNA-seq (A, B), CapSeq (C, D) and small RNA-seq (E) in *cuff* mutants, and *cuff* mutants expressing either *mel*-*cuff* or *sim*-*cuff.* Diagonal represents x=y. Points in red show y/x>3. p value for differences obtained by Wilcoxon test. (F) Genome browser view of 42AB piRNA cluster shows long RNA-seq profiles of WT control, *cuff* mutants, and *cuff* mutants expressing either *mel*-*cuff* or *sim*-*cuff*. Both *cuff* mutants and *sim-cuff* expressed in *cuff* mutants lead to splicing of precursor transcripts. (G) Bar graphs showing ChIP vs. Input enrichment for Rhi at the indicated heterochromatic repeats in *cuff* mutant ovaries expressing *mel-cuff* (black) or *sim-cuff* (grey).

**Figure S2, related to figure 2:**
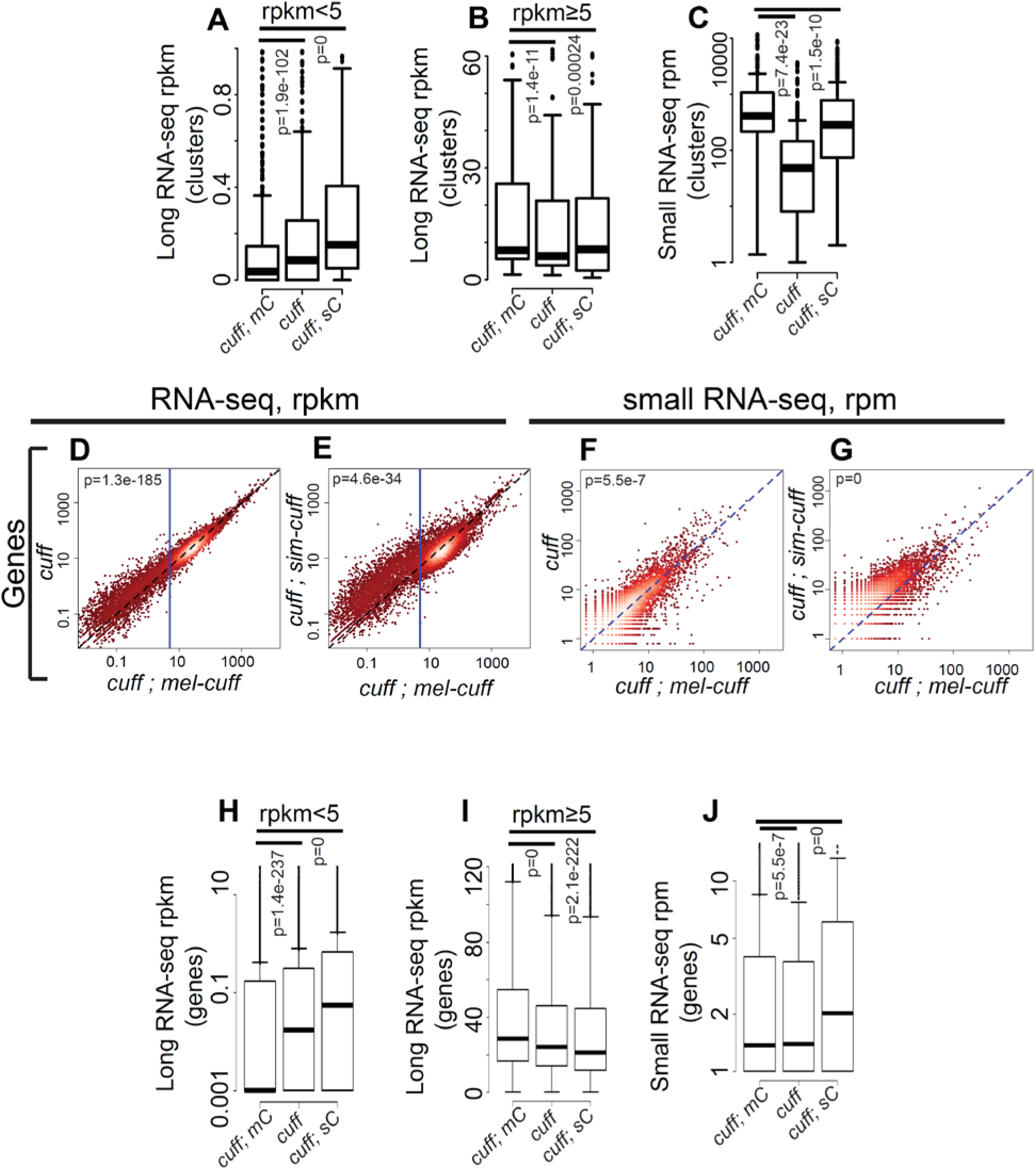
(A-C) Boxplots showing comparisons of RNA-seq signal (A and B) and small RNA-seq signal (C) at piRNA clusters in ovaries with genotypes *cuff* mutant, *cuff* mutant expressing either *mel-cuff* or *sim-cuff*. (A) and (B) are for rpkm<5 and rpkm>5 in *cuff* mutant expressing *mel-cuff* respectively. p values obtained by Wilcoxon test. (D-J) Boxplots and scatterplots showing gene mapping long-RNA-seq (D, E, H and I) and small RNA-seq (F, G and J) in *cuff* mutants, and *cuff* mutants expressing either *mel*-*cuff* or *sim*-*cuff.* Diagonal represents x=y. Points in red show y/x>3. p value for differences obtained by Wilcoxon test. Blue vertical lines in (D) and (E) show x=5. (H) and (I) are for rpkm<5 and rpkm>5 in *cuff* mutant expressing *mel-cuff* respectively.

**Figure S3, related to figure 3:**
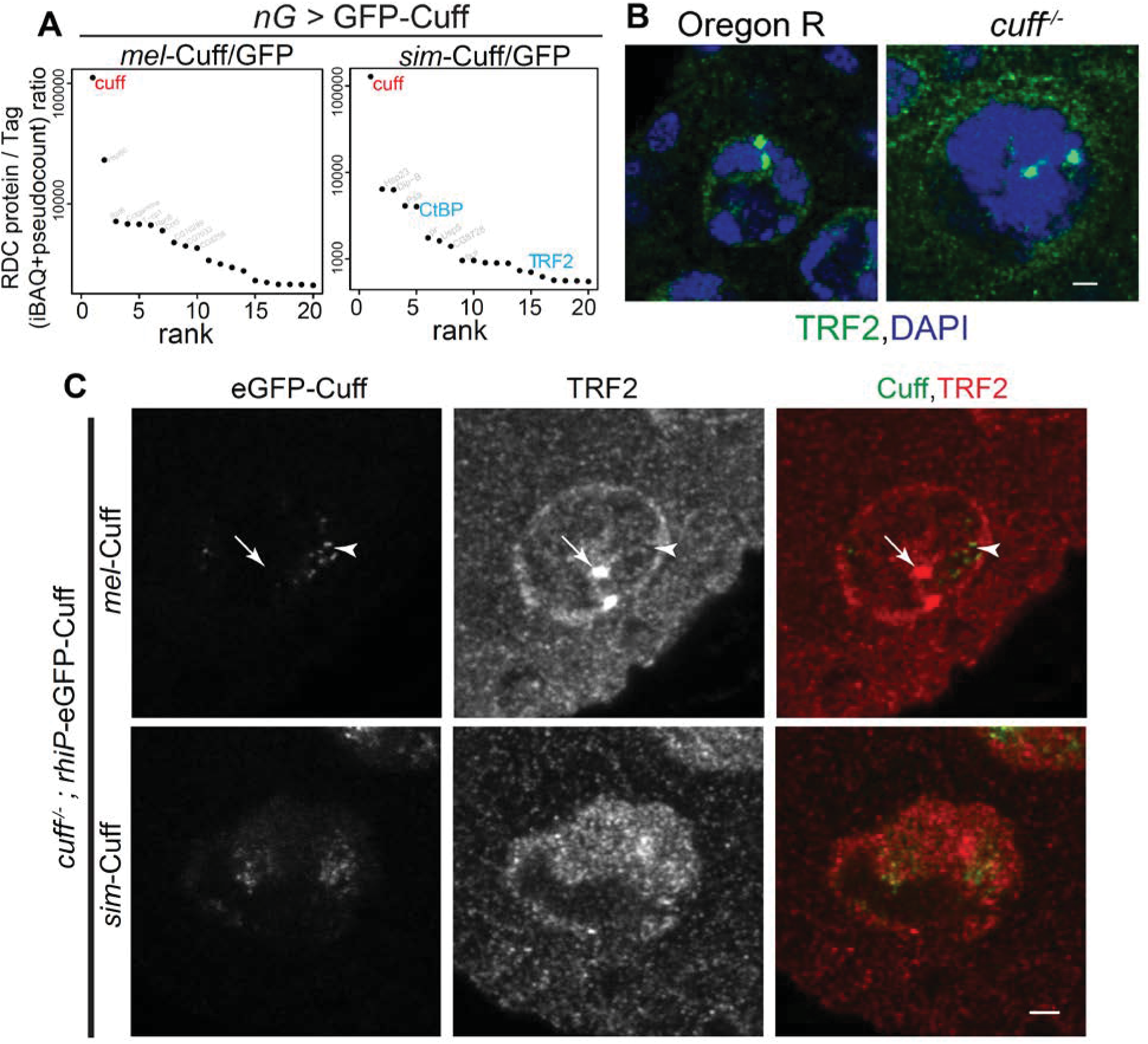
(A) Mass spectrometric analysis of over-expressed *nanos*-Gal4 driven *mel*-Cuff and *sim*-Cuff binding proteins. Graphs show ratios of iBAQ value of a bound protein in a RDC protein IP vs. tag control IP ranked by ratio values. Cuff is shown in red, TRF2 and CtBP in blue. (B) Localization of TRF2 with respect to DNA (DAPI) in the germline nuclei of WT control (Oregon R) and *cuff* mutant. TRF2 and DNA (DAPI) are shown in green and blue respectively. Scale bar, 2 µm. (C) Localization of GFP tagged Cuff with respect to TRF2 in the germline nuclei of *cuff* mutants expressing *rhi* promoter driven *mel*-Cuff or *sim*-Cuff. Color assignments for merged images shown on top. Scale bar, 2 µm. Arrowheads and arrows denote locations of *mel*-Cuff or TRF2 foci respectively.

**Figure S4, related to figure 3:**
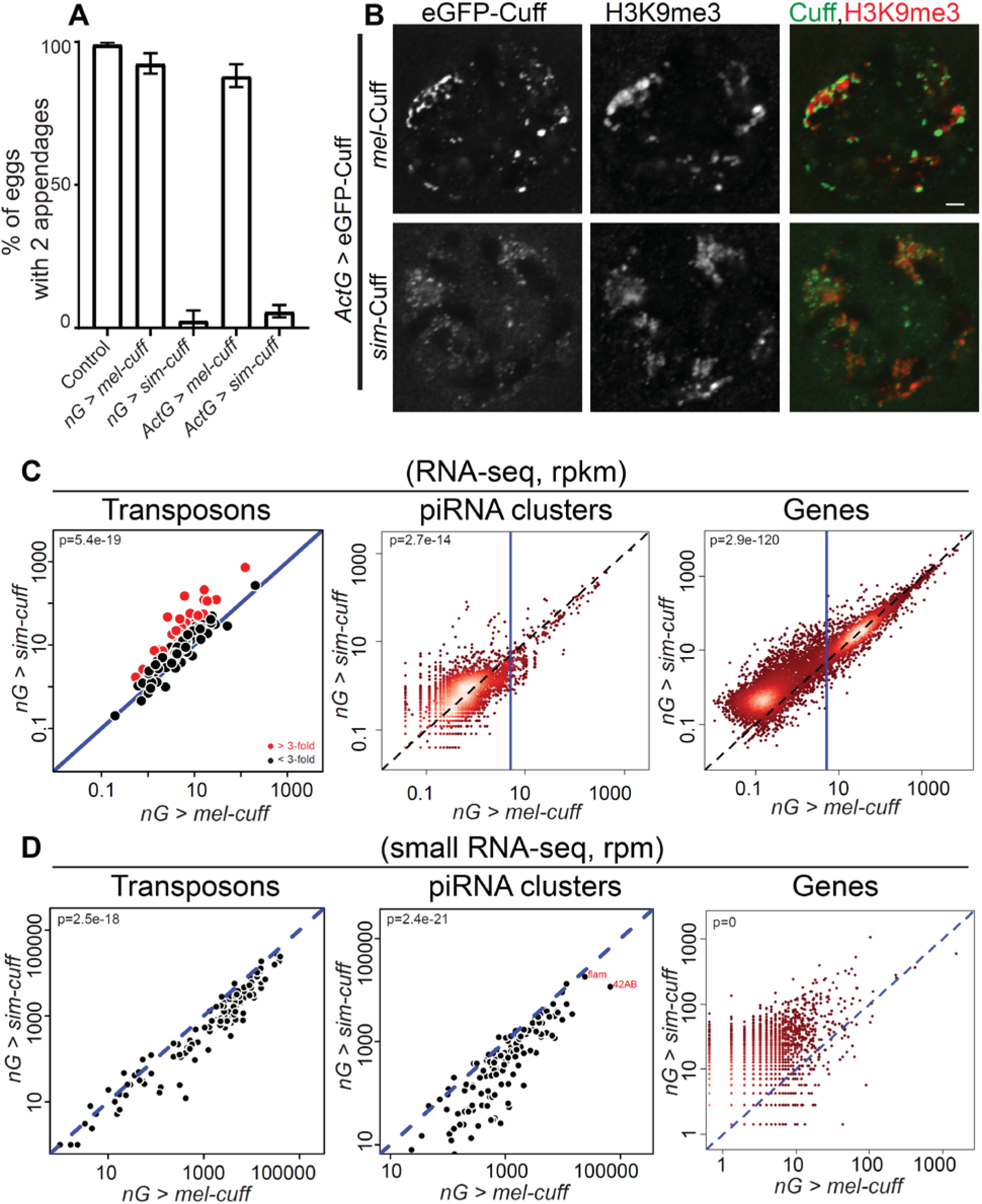
(A) Bar graphs showing percentages of eggs with 2 appendages produced by control (*w*^1^; *Sp*/CyO), flies over-expressing either *mel*-*cuff* or *sim*-*cuff* by either *nanos*-Gal4 (nG) or *Act5C*-Gal4 (*Act*-Gal4) drivers. Error bars show standard deviation of three biological replicates, with a minimum of 200 embryos scored per replicate, except for *nanos*-Gal4 driven *sim-cuff* where average of 50 eggs were scored. (B) Localization of GFP tagged Cuff with respect to H3K9me3 marked chromatin in germline nuclei of *Act*-Gal4 driven *mel*-Cuff or *sim*-Cuff. Color assignments for merged images shown on top. Scale bar, 2 µm. (C and D) Scatterplots showing comparisons of RNA-seq signal (C) and small RNA-seq signal (D) at transposons, piRNA clusters and genes in *nanos*-Gal4 driven *sim-cuff* vs. *nanos*-Gal4 driven *mel-cuff.* Diagonal represents x=y. Red points in (C, transposons plot) show y/x>3. Each point in (C, piRNA clusters plot) shows 1kb bin of a piRNA cluster. p value for differences obtained by Wilcoxon test.

**Figure S5, related to figure 4:**
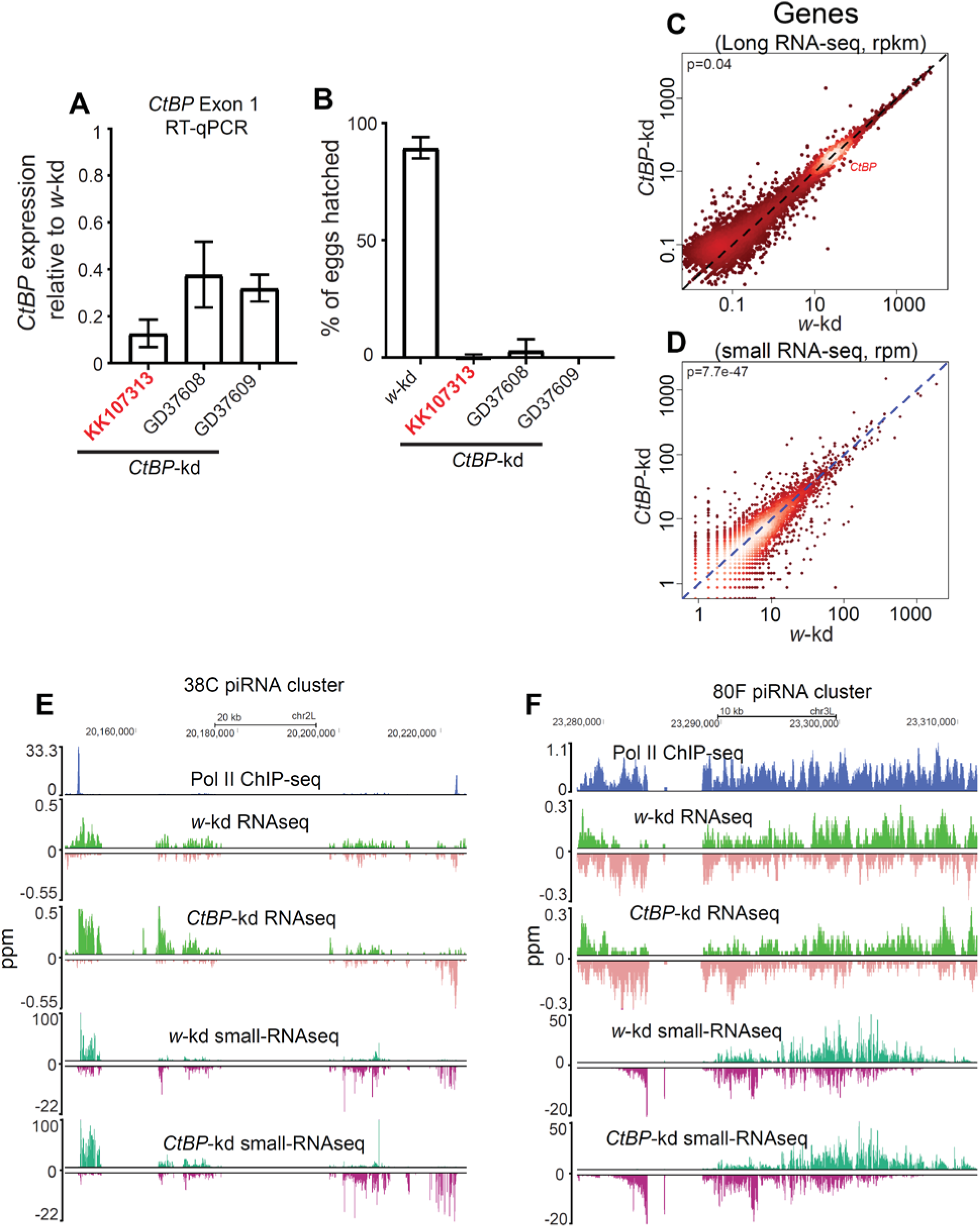
(A) Bar graphs showing *CtBP* exon-1 expression in ovaries of different *CtBP*-kd lines vs. *w*-kd line by RT-qPCR. Error bars show standard deviation from 3 biological replicates. (B) Bar graphs showing percentages of hatched eggs produced by flies with germline knock-down by different *CtBP*-kd lines. Error bars show standard deviation of three biological replicates, with a minimum of 200 embryos scored per replicate. (C, D) Scatterplots showing comparisons of RNA-seq (C) and small RNA-seq signal at genes in *CtBP-*kd vs. *w-*kd ovaries. Each point on the scatterplots shows rpkm or rpm value for a gene. Diagonal represents x=y. p value for differences obtained by Wilcoxon test. (E, F) Genome browser view of RNA-seq (top) and small RNA-seq (bottom) profiles at 38C (E) or 80F (F) piRNA clusters from *w-*kd and *CtBP-*kd ovaries. Pol II ChIP-seq peaks in *nanos*-Gal4 driven *mel*-Cuff ovaries are shown (in blue) to denote cluster promoters at 38C. Such distinct promoter is absent at 80F cluster.

**Figure S6, related to figure 6:**
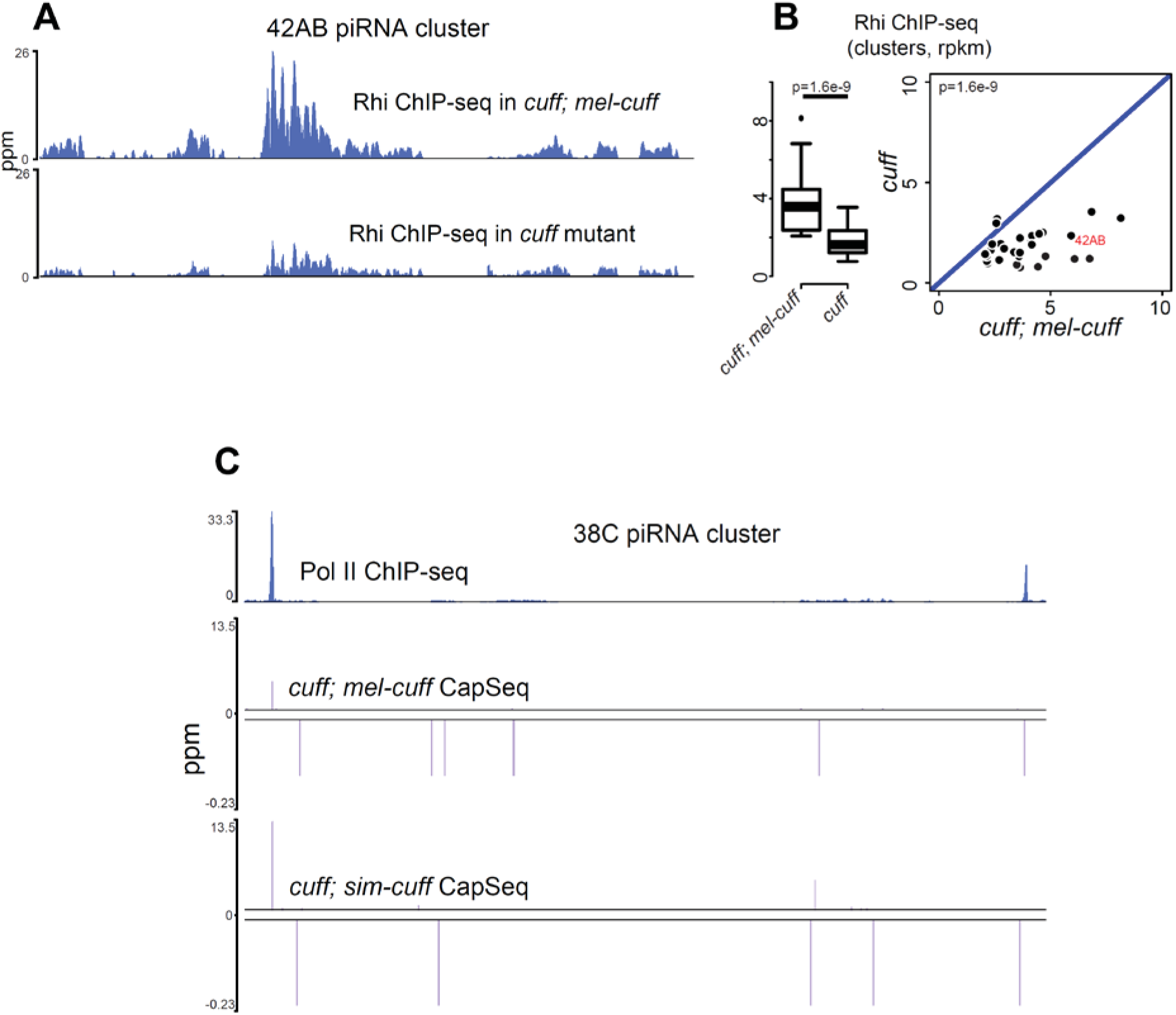
(A) Genome browser view of Rhi ChIP-seq profiles at 42AB piRNA cluster in the ovaries of *cuff* mutant and *cuff* mutant expressing *mel-cuff*. (B) Boxplot and scatterplot showing comparisons of ChIP/Input values for Rhi at piRNA clusters in ovaries with genotypes *cuff* mutant vs. *cuff* mutant expressing *mel-cuff*. The clusters with prominent Cuff or Rhi binding (rpkm>2) in *cuff* mutant with *mel-cuff* control were used for analysis. Diagonal represents x=y. p value for differences obtained by Wilcoxon test. (C) Genome browser view at 38C cluster, showing Pol II ChIP-seq in *nos*-Gal4 driven *mel-cuff* and CapSeq signals in *cuff* mutants expressing either *mel-cuff* or *sim-cuff*.

